# Isolation and characterization of diverse *Veillonella* phages reveal novel lineages and distinct host interactions

**DOI:** 10.64898/2026.11.01.763356

**Authors:** Anubhav Basu, Hyelan Lee, Jackie Fenner, Junhan Ruan, Christopher A. Gaulke, Carin K. Vanderpool

## Abstract

The human oral microbiome is diverse and home to many species of bacteria, archaea, microeukaryotes, fungi and viruses. Homeostasis in the oral microbiome is important for health and dysbiosis in these communities leads to periodontal and systemic diseases. *Veillonella* are gram-negative, anaerobic, lactate-fermenting early colonizers of the mouth and are abundant in healthy oral microbiomes. Although considered commensals, *Veillonella* serve as bridging species in dental biofilms, promoting later colonization by pathogens, with mounting evidence supporting their roles as accessory pathogens in dental caries and periodontal disease. Bacteriophages are key regulators of bacterial community composition and are important drivers of microbial diversity, evolution, and ecology of the oral environment. Despite the ecological importance of *Veillonella* in the oral microbiome, phages infecting *Veillonella* remain largely uncharacterized. Here, we report the isolation of ten phages infecting natural isolates of three *Veillonella* species, representing seven distinct phage genomes. These phages exhibit diverse morphologies, genome organizations, and host ranges, with infection restricted to strains within the species of isolation. Several phages establish stable lysogeny, confer resistance to superinfection, and differ substantially in spontaneous production of infectious phage progeny. Genetic analysis of resistant mutants implicates lipopolysaccharide O-antigen as the receptor for one phage. Although none of the isolated phages had close matches in existing oral virome catalogs, related sequences were detected in oral metagenomes from geographically diverse populations. Together, these findings establish an experimentally characterized collection of *Veillonella* phages and connect previously uncharacterized oral viral diversity to bacterial hosts and infection phenotypes.

**Importance:** Many bacteria in the mouth form structured biofilms on teeth and gum tissue. Some bacteria, including *Veillonella*, help pathogenic bacteria establish a foothold in these communities, contributing to cavities and periodontal disease. Viruses that infect bacteria, called bacteriophages, naturally regulate bacterial populations, but almost nothing is known about phages that infect *Veillonella* species. This study identifies and characterizes novel *Veillonella* phages genetically and functionally, revealing diversity in their structure, host range, and behavior. Understanding their biology opens the door to studying how these phages impact their hosts in natural environments as well as other members of the oral community.

## Introduction

The human oral microbiome is home to a diverse collection of microorganisms including bacteria, archaea, fungi, microeukaryotes and viruses that represents the second most diverse microbiota in the body (1). It harbors more than 700 bacterial species including members of the Actinomycetota, Bacteroidota, Bacillota, Fusobacteriota, Pseudomonadota, Saccharibacteria and Spirochaetota (2, 3). The microbiome exists in the form of a highly complex and structured biofilm and plays an important role in maintaining homeostasis and protecting the oral cavity from pathogens (3). Indeed, a dysbiotic oral microbiota is associated with development of diseases such as dental caries, periodontitis, and oral cancers (3–5). Further, there is increasing evidence that a dysbiotic oral microbiota can lead to extra-oral diseases like Alzheimer’s, colorectal cancer and cardiovascular disease, among others (5–8).

*Veillonella* are anaerobic, gram-negative early colonizers of the mouth and are abundant in human oral microbiomes (9, 10). The genus comprises 16 characterized species of which eight are commonly found in the oral cavity (10) where they play an important role in formation of multispecies biofilms (11, 12). For example, they bind to surface receptors of early colonizing streptococci and themselves act as binding partners for late colonizing pathogens (13–16). *Veillonella* also participate in metabolic interactions as exemplified by their relationship with oral streptococci where lactate produced by streptococci is consumed by *Veillonella* and converted into acetate and propionate (17). Beyond merely acting as a binding partner, *Veillonella* can also modulate the biofilm microenvironment; for example, catalase produced by some *Veillonella* species can detoxify hydrogen peroxide produced by streptococci (18) and they produce heme that can be used as a nutrient by other members of the community (20). These activities may promote the proliferation of less aerotolerant pathogenic bacteria and so *Veillonella* have been proposed as ‘accessory pathogens’ (10). In dental caries, *Veillonella* abundance tracks that of acid-producing species, and among children with no previous caries it predicted future disease (19), with *V. parvula* among the species most consistently recovered from severe early childhood caries (21). Further, *V. parvula* produces a soluble signal that supports the growth of the pathogen *Porphyromonas gingivalis* at low cell densities (22) and in diabetes associated periodontitis, nitrate generated by inflamed gingival epithelium enhances *P. gingivalis* growth and virulence in a *V. parvula* dependent manner (23). Thus, the apparent participation of *Veillonella* in multiple oral diseases processes and interactions with key players in those disease processes make them an attractive indirect target for microbiome-based interventions.

The oral cavity contains 10^8^ virus-like particles per milliliter of saliva, with the majority being bacteriophages (24, 25). Phages play an important role in modulating the oral microbiome and in turn modulate oral health (26). Phages infecting members belonging to major oral phyla have been isolated or predicted from metagenomes (27, 28). Recently, an effort to catalog oral phages resulted in a collection of more than 180,000 phage genomes with more than half of the classified phage clusters representing lytic phages (28). Even where phages haven’t been isolated, their genomic footprints reveal they are active players in oral microbial evolution and virulence. A recent study identified prophages in the genomes of periodontal pathogens and found that prophages carry putative genes for horizontal gene transfer, modulation of host metabolism and anti-host defense systems such as anti-CRISPRs (26).

Despite the importance of *Veillonella* in the oral microbiome, phages infecting them remain understudied. Only one prior study has reported the isolation of 25 *Veillonella* phages and separated them into two distinct groups based on plaque morphology and serological characteristics, suggesting that diversity exists among *Veillonella* phages (29). Another study used the same phages to classify 21 *Veillonella* isolates into 4 groups and reported basic growth characteristics of a few phages (30). However, these studies predate the genomic era, and no genomic, lifestyle or ecological characterization was performed. By contrast, phages infecting other abundant oral genera such as *Streptococcus* (31, 32), *Fusobacterium* (33), and *Actinomyces* (34) have received substantially more attention, making *Veillonella* an outlier in oral phage research.

Here we describe the isolation of novel phages infecting natural isolates of three different *Veillonella* species. The phages are diverse and have different morphologies, host ranges, lifestyles and gene content. Comparison to known phages using genome similarity suggested that they represent distinct, novel genera. Further, we show that several of these phages are temperate and can integrate into their host genomes as prophages and provide superinfection immunity. Some prophages spontaneously induced at high levels during exponential growth of their host while others fail to induce even under stress conditions. We show that the isolated phages are absent from existing catalogues, further underscoring their novelty but related phages are present in different geographical areas with varying abundance. This study presents the first genomic and functional characterization of naturally occurring *Veillonella* phages, revealing unexpected diversity in phage lifestyle and host interaction strategies.

## Results

### Isolation of *Veillonella* strains from natural environments

We were broadly interested in identifying diverse *Veillonella* strains and phages from the oral microbiome and studying interactions between phage-host pairs. For this, we first isolated *Veillonella* strains from human oral rinse samples by plating on selective media and using primers specific for the *rpoB* gene. We isolated a total of nine strains, including two strains of *V. parvula* (ABVpa1-2), five of *V*. *atypica* (ABVat1*-5*), and two of *V. nakazawae* (ABVna1-2) (Table S1). We also included the type strain *V. parvula* ATCC 10790 (hereafter referred to as ATCC 10790) in our study. We sequenced the isolates using long read sequencing and placed the assemblies within the GTDB reference tree to understand their phylogenetic relationship (Fig. 1A). Among the *V. parvula* strains, ABVpa1 and ABVpa2 were nearly identical, sharing 100% average nucleotide identity (ANI) and differing by only 9 SNPs and 4 small indels across their ∼2.1 Mb genomes (Table S2, Fig. S1). Both clustered together with the GTDB reference genome *V. parvula* G (GCA_032550495.1) and formed a clade separate from ATCC 10790, which shared 95.9% ANI with the two isolates. The *V. atypica* strains clustered together, with ABVat4 being closest to the reference strain *ATCC 17744*. Strains ABVat1 and ABVat3 were almost identical, sharing 100% ANI and differing by only 4 small indels with no SNPs, while ABVat5 was the most divergent from the reference strain, sharing ∼96.5% ANI with the other *V. atypica* isolates (Fig. S2). ABVat5 was the only isolate that harbored a plasmid (Table S1). The two *V. nakazawae* strains, ABVna1 and ABVna2, clustered together with reference strain *V. nakazawae* T1-7 but were on a separate branch, sharing 95.5% ANI with each other and 84-87% ANI with strains of the other two species, consistent with their classification as a distinct species.

**Figure 1.**
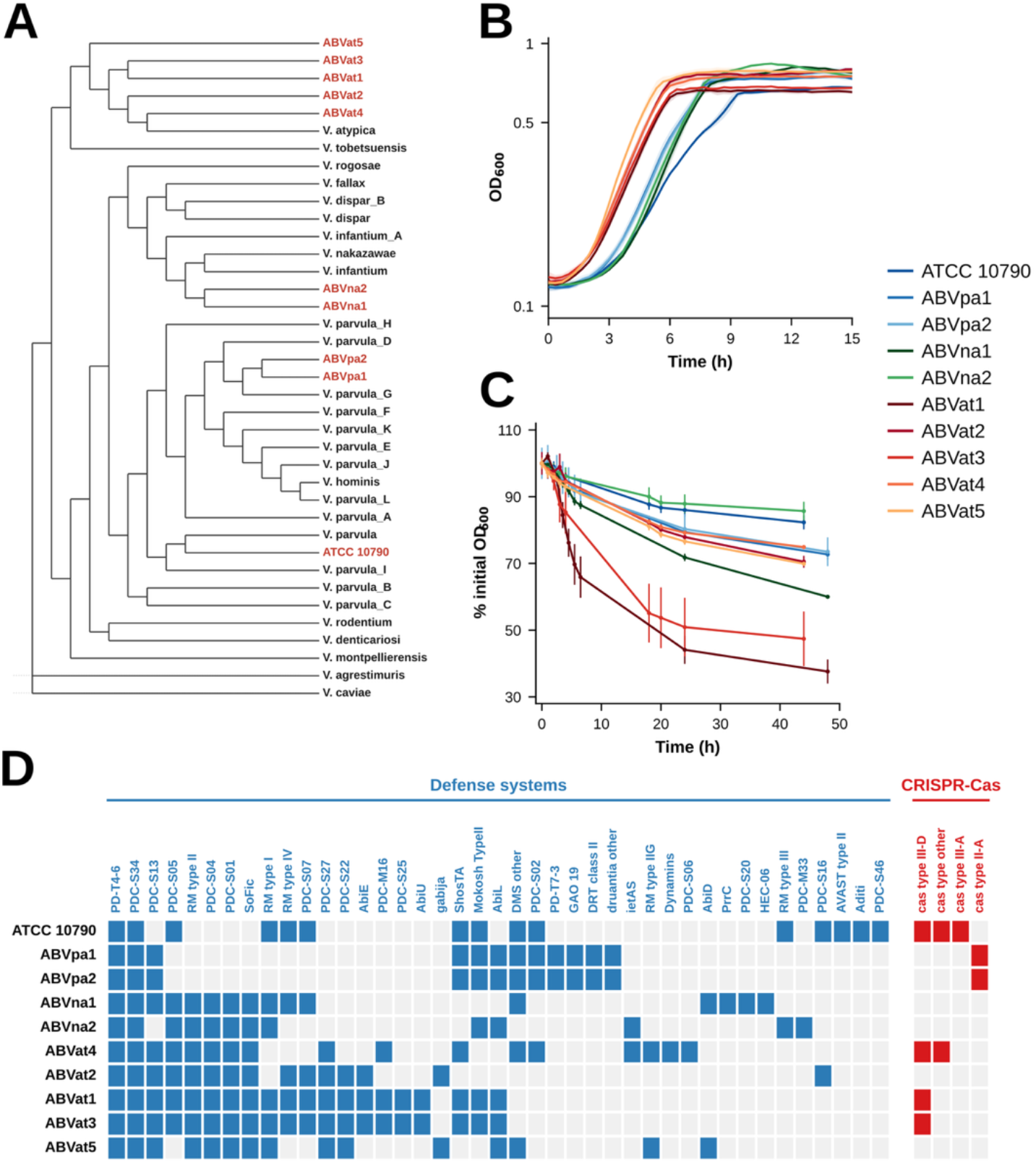
Isolation and characterization of novel *Veillonella* strains. (A) Cladogram constructed using GTDB-Tk, showing the placement of the novel isolates and the type strain ATCC 10790 within the broader *Veillonella* genus. Isolates from this study are highlighted in red. Branch lengths are not to scale. (B) Growth curves of all ten *Veillonella* strains in rich SK media. Optical density (OD600) was measured at 20 minute intervals over 15 hours. Lines represent means of n=3 biological replicates and shading indicates standard deviation. (C) Autoaggregation of novel *Veillonella* isolates as measured by a sedimentation assay. Values represent the percentage of initial OD600 remaining in suspension over time in a cuvette. Points represent means of n=3 biological replicates and error bars indicate standard deviation. (D) Annotated phage defense systems of all ten strains identified using PADLOC. Blue tiles indicate the presence of a defense system, red tiles indicate presence of CRISPR-Cas systems and grey tiles indicate absence of the defense system.

To further characterize the novel isolates, we measured their growth in rich media. The *V. atypica* isolates grew fastest (Td = 1.35-1.70 h) and ATCC 10790 slowest (Td = 2.21 h), while the *V. nakazawae* strains had intermediate doubling times but the longest lag phases at 3.1-3.3 h against 1.6-1.8 h for the *V. atypica* strains (Fig. 1B, Table S3).

Many oral bacteria, including *Veillonella,* aggregate either with themselves (autoaggregation) or with other species (coaggregation) (14). To measure autoaggregation in our isolates, we performed a sedimentation assay which measures how quickly cells clump together and settle at the bottom of a cuvette (Fig. 1C). The *V. atypica* isolates aggregated more than the other species with ABVat1 showing the strongest phenotype with almost 60% reduction from the initial OD after 24 hours while strains ATCC 10790 and ABVna2 autoaggregated the least and had an OD reduction of only 15% after 48 hours.

The nature of phage-bacteria interactions is influenced by several traits of both the phage and its host. Among these traits is the array of anti-phage defenses that the host possesses. Using the Prokaryotic Antiviral Defence LOCator (PADLOC), we identified a diverse array of phage defense genes in our isolates (Fig. 1D). We found CRISPR-Cas systems were present in all *V. parvula* and some *atypica* isolates, but absent from the *V. nakazawae* isolates. Also, restriction–modification (RM) systems were present in every isolate with RM type II being the most abundant among these strains.

### Isolation of novel *Veillonella* phages

Using the panel of novel *Veillonella* isolates as hosts, we used classical enrichment techniques to isolate 10 novel phages. The phages collectively infect at least one representative of each of the three *Veillonella* species (Table S4). In total, three phages infecting *V. parvula*, six infecting *V. atypica* and one phage infecting *V. nakazawae* were isolated. Phages were named using the vB nomenclature system, in which the host genus initial (V - *Veillonella*), species abbreviation (pa - *parvula*, at - *atypica*, na - *nakazawae*), and morphology code (P - podovirus, M - myovirus and S - siphovirus) are encoded in the name, followed by a sequential number (P1 - phage 1, P2 - phage 2 etc.).

To determine the morphology of the isolated phages we performed transmission electron microscopy (Fig. 2A). All three phages infecting *V. parvula* (vB_VpaP_P1, vB_VpaP_P2 and vB_VpaP_P3) and two phages infecting *V. atypica* (vB_VatP_P4 and vB_VatP_P5) were found to be podophages. Of the two isolated myophages, one infected *V. atypica* (vB_VatM_P1) and the other *V. nakazawae* (vB_VnaM_P1), while the lone siphophage infected *V. atypica* (vB_VatS_P6). The podophages had a 59.8 ± 2.7 nm capsid with a short tail. Among the myophages, vB_VatM_P1 had a bigger capsid and tail (68.2 ± 1.9 nm and 133 ± 2.3 nm respectively) compared to vB_VnaM_P1 (47.7 ± 1.3 nm and 76.6 ± 1.7 nm respectively). The siphophage had a small (46.8 ± 1.2 nm) capsid and a long (169.4 ± 7.2 nm) flexible tail (Fig. S2).

**Figure 2.**
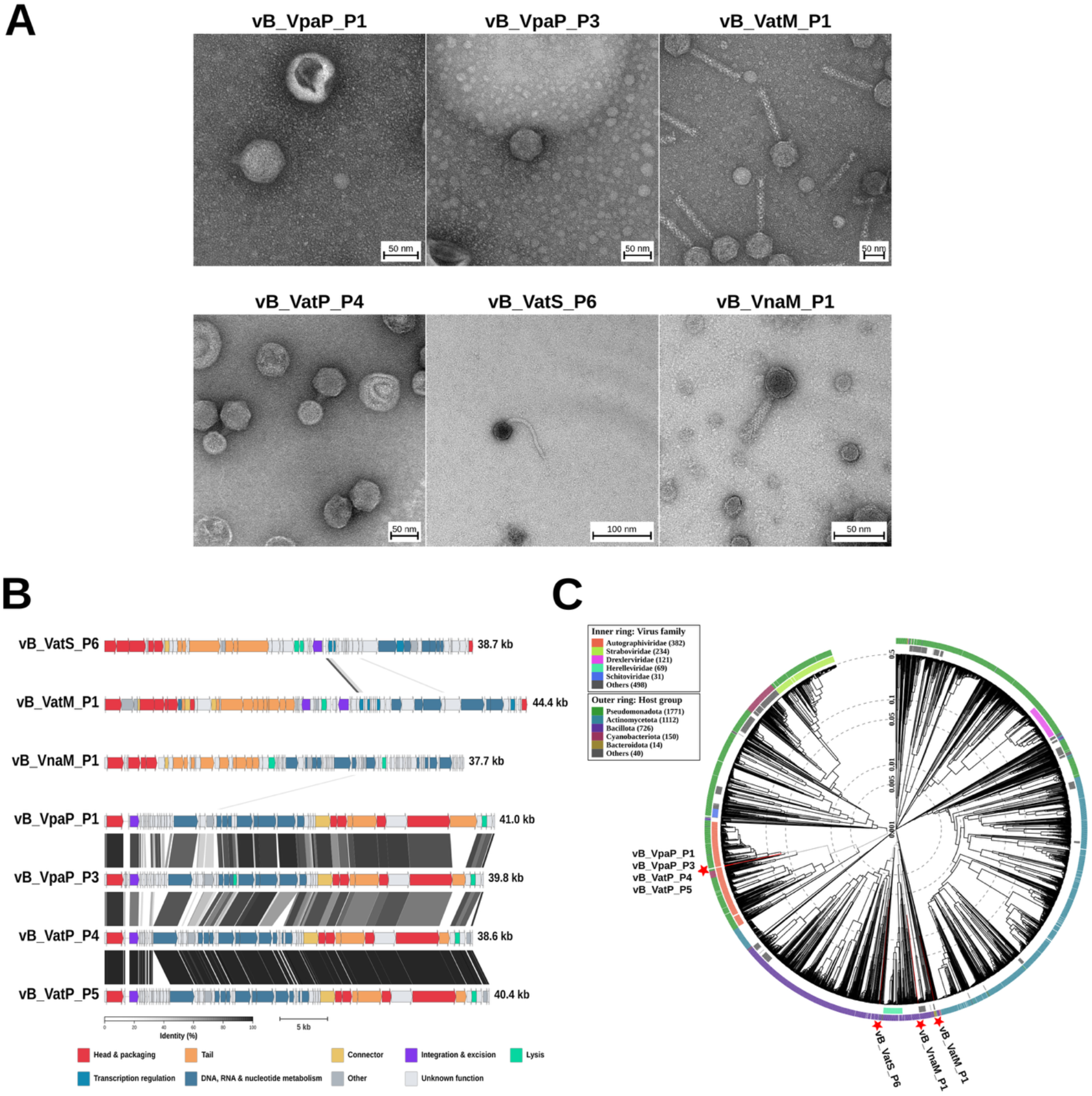
Genomic characterization and phylogenetic analysis of novel *Veillonella* phages. (A) Representative transmission electron microscopy (TEM) images of the novel *Veillonella* phages. Scale bars are present in the insets. (B) Pharokka annotated genome maps of the novel phages, drawn to scale. Arrows represent coding sequences colored by functional category as indicated in the legend. Synteny among the genomes, generated with CLINKER, is shown as a heatmap ranging from 30% (white) to 100% (black) identity. (C) Phylogenetic tree of the novel phages generated with ViPTree, placing them among reference genomes. Phages from this study are marked with stars. Host phyla and phage family, wherever known, are indicated in the outer circular rings. Inner dashed circular rings indicate genome similarity as measure using Sg.

Sequencing revealed that all isolated phages had double-stranded DNA genomes ranging from 37 kb - 44 kb (Fig. 2B, Table S4). We found that phages vB_VatM_P2 and vB_VatM_P3 differed from vB_VatM_P1 by only one and two SNPs respectively, while phages vB_VpaP_P1 and vB_VpaP_P2 were also similar to each other with a small number of SNPs (Table S5). Hence, only vB_VatM_P1 and vB_VpaP_P1 were selected as representatives for further evaluation. Curiously, phages vB_VatP_P4 and vB_VatP_P5 were identical except for a cluster of eight genes present in vB_VatP_P5. All phage genomes assembled as circular contigs. However, analysis of sequencing reads from two podophages, vB_VpaP_P1 and vB_VpaP_P3, revealed evidence of linear genomes with short direct terminal repeats, consistent with a defined-terminus DNA packaging strategy (Fig. S3). The genome termini and packaging strategies of the remaining phages could not be determined confidently from the available sequencing data.

Synteny analysis revealed broad conservation of gene order among all the podophages, while the myophages and siphophage had essentially no synteny between each other or with the podophages (Fig. 2B). To determine how these isolated phages relate to characterized phages, we compared their genomes against the Virus-Host DB reference database using ViPTree (35) which quantifies similarity between phage genomes with Sg (36), a normalized tBLASTx based score ranging from 0 (no detectable similarity) to 1 (identical). We found that none of the isolated phages were meaningfully similar to any of the reference genomes with the highest Sg observed for any pairing being 0.146 between vB_VatP_P4 and *Yersinia* phage YpP-G, underscoring the novelty of these phages (Fig. 2C, Table S6). Despite this general lack of close relatives, the podophages all had their nearest hits (Sg = 0.13 - 0.15) to members of the family *Autographiviridae*, a group of phages defined by phage encoded single subunit RNA polymerase (37). Further, all isolated podophages contain such a single subunit RNA polymerase. This suggests that the podophages may represent one or more novel genera within the *Autographiviridae*. In contrast, the myophages and siphophage showed no consistent family level signal among their nearest hits, which spanned phages of taxonomically unrelated hosts including *Vibrio*, *Faecalibacterium*, *Priestia*, and *Lactobacillus* (Sg ≤ 0.10, 0.04, and 0.04, respectively, Table S6), consistent with these phages representing highly divergent lineages with no clear relationship to established families.

We found at least one integrase gene within each phage genome (Fig. 2B) except vB_VnaM_P1, which lacks an annotated integrase which is consistent with a strictly lytic lifestyle. Further, we found that the phages encoded distinct complements of genes involved in transcription, DNA replication, nucleotide metabolism, host-defense evasion, recombination, and lysis (Table S7). On average, 58% of predicted coding sequences across the phage genomes lack functional annotation, underscoring their novelty and the remarkable diversity of phage gene products.

### Novel phages are species restricted but show some flexibility in host range

Most of the isolated phages formed turbid plaques on their host lawns, consistent with a presumed temperate lifestyle and ability to form lysogens. However, the turbidity of plaques varied between the phages (Fig. 3A). The siphophage vB_VatS_P6 made highly turbid plaques which were difficult to see while the putative strictly lytic phage vB_VnaM_P1 made large, clear plaques. The other phages made plaques of intermediate turbidity. Interestingly, the two podophages that infect ATCC 10790 had distinct plaque morphologies: vB_VpaP_P3 made clearer plaques while vB_VpaP_P1 made more turbid plaques.

**Figure 3.**
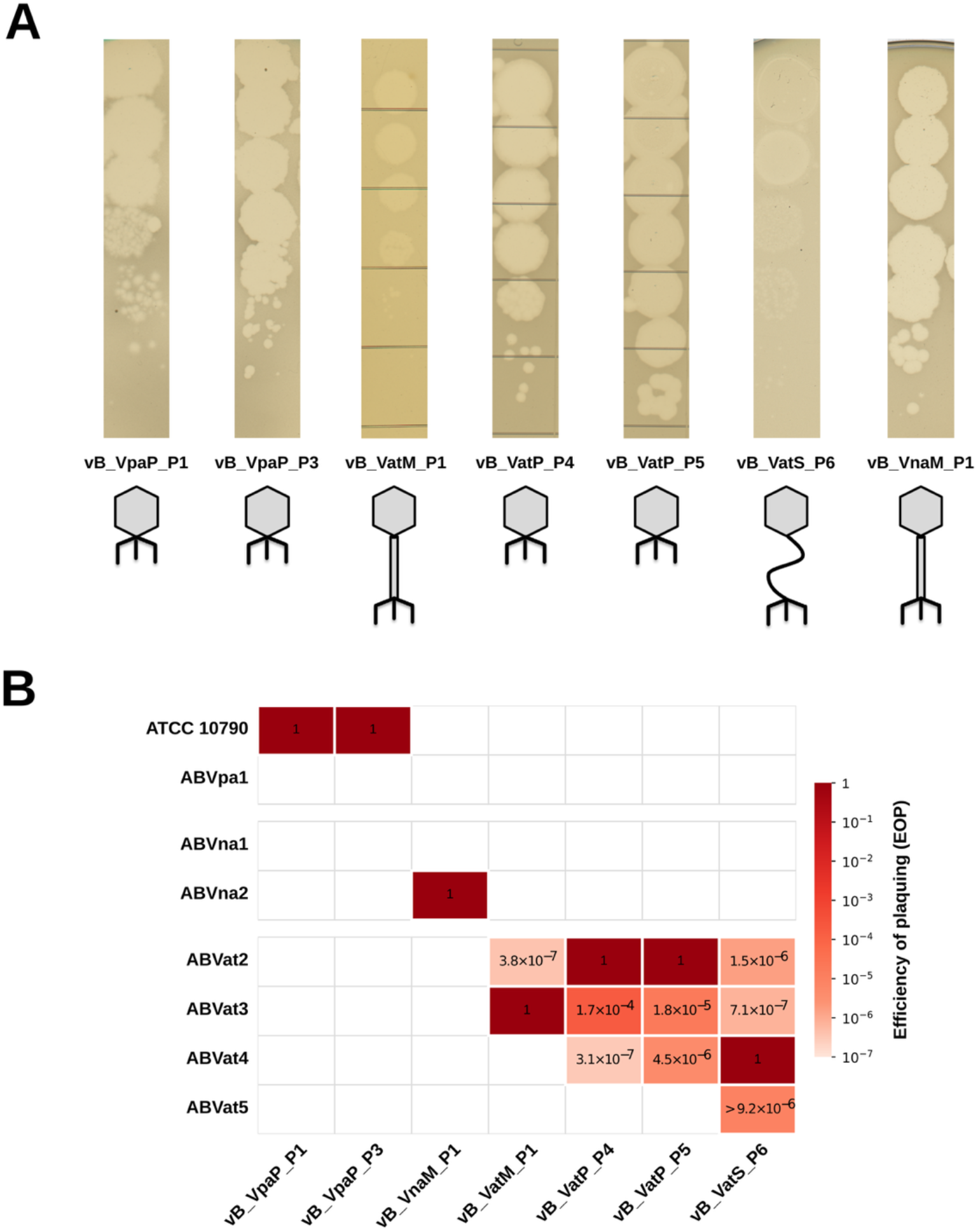
Infection characteristics and host range of novel *Veillonella* phages. (A) Plaque morphologies of the novel phages on their isolation host. Phage morphology (Podophage, Myophage or Siphophage) is indicated by cartoons. (B) Host range of the isolated phages shown by Efficiency of Plaquing (EOP) on a strain relative to its isolation strain. Actual EOP values are indicated in the tiles of the heatmap and are derived from n=3 biological replicates. White tiles represent no infection.

Next, we tested the host range of the seven unique phage genomes against our panel of *Veillonella* isolates using the efficiency of plaquing (EOP) relative to the isolation host (Fig. 3B). ABVpa2 and ABVat1 were omitted because each is identical by ANI to an isolate already in the panel (Fig. S1). We saw that infection was strictly species restricted, and no phage produced plaques on a host outside its species of isolation. Within *V. parvula* and *V. nakazawae*, host range was limited to the isolation host alone and the phages failed to infect other strains of these species. On the other hand, the *V. atypica* phages infected multiple strains within the species, but at reduced efficiency outside the isolation host. The siphophage vB_VatS_P6, isolated on ABVat4, had the broadest host range and plaqued on all other *V. atypica* strains. The podophages vB_VatP_P4 and vB_VatP_P5, both isolated on ABVat2, plaqued on ABVat3 (EOP 1.7 × 10⁻⁴ and 1.8 × 10⁻⁵) and on ABVat4 (3.1 × 10⁻⁷ and 4.5 × 10⁻⁶), but not on ABVat5. Myophage vB_VatM_P1, isolated on ABVat3, had the narrowest range, plaquing on ABVat2 alone and at the lowest efficiency we recorded (3.8 × 10⁻⁷). This pattern of infectivity could not be explained by genome similarity alone. For example, strains ABVat3 and ABVat4 are equally similar to ABVat2 (97.0% ANI to both), yet vB_VatP_P4 plaqued approximately 570-fold more efficiently on ABVat3 than on ABVat4.

### Novel *Veillonella* phages readily form lysogens *in vitro*

To determine whether the temperate phages could integrate into their hosts’ chromosomes and establish stable lysogeny, we performed an efficiency of lysogeny (EOL) assay for each phage-host pair, in which serial dilutions of the host were plated on phage-seeded and non-seeded control plates and the fraction of surviving colonies was determined (41). Survival frequencies varied dramatically across the five phages tested, spanning nearly three orders of magnitude (Fig. 4A). The highest survival frequency was observed for vB_VatS_P6 on its host ABVat4 (90.19 ± 4.41%), while vB_VatM_P1 had the lowest (0.20 ± 0.08%). Intermediate survival frequencies were observed for the podophages vB_VpaP_P1 (12.61 ± 4.5%), vB_VatP_P5 (10.56 ± 4.67%), and vB_VpaP_P3 (1.85 ± 0.77%). Interestingly, vB_VpaP_P1 and vB_VpaP_P3 which infect the same host strain, differed approximately 7-fold in survival frequency, suggesting that phage-specific properties influence the outcome of infection.

**Figure 4.**
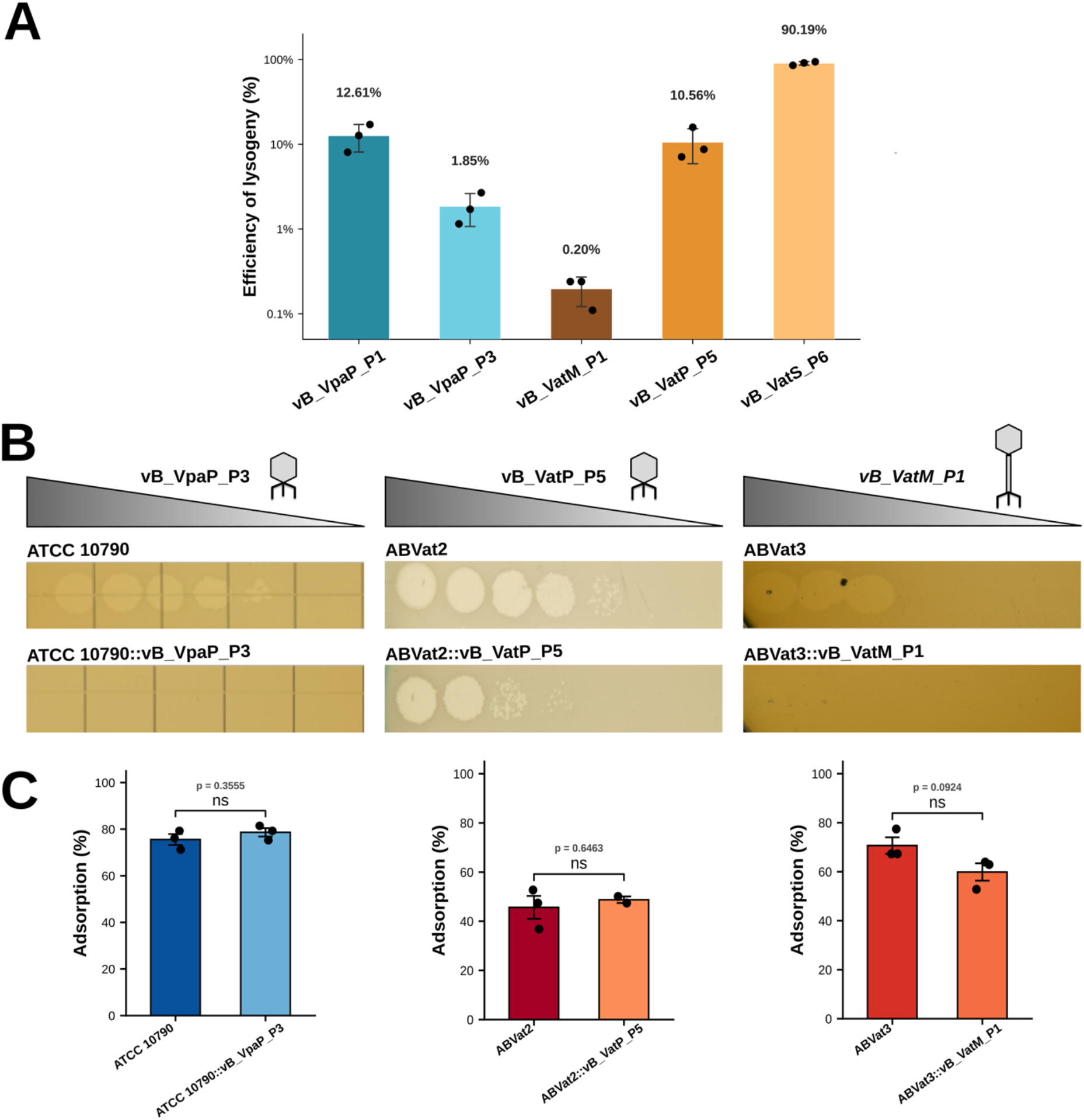
Lysogeny, superinfection immunity, and adsorption in *Veillonella* phage-host pairs. (A) Efficiency of lysogeny (EOL) assay for five phages on their respective hosts, showing the fraction of colonies surviving on phage-seeded plates relative to non-seeded control plates. Survival was used as a proxy for lysogeny, which was independently confirmed by genome sequencing for three representative phage-host pairs. Bars show mean ± SD of n=3 biological replicates. (B) Superinfection immunity displayed by lysogenized strains. Phages spotted onto lawns of *ATCC 10790::vB_VpaP_P3 and ABVat3::vB_VatM_P1 fail to form plaques, while ABVat2::vB_VatP_P5 exhibits partial resistance to superinfection.* The same phages spotted onto lawns of the susceptible wild type *hosts* readily form plaques. (C) Adsorption assays comparing phage adsorption fraction of wild type and lysogenic strains for each of the three pairs in (B). Bars show mean ± SD of n=3 biological replicates. Statistical comparisons between wild type and lysogen were performed using unpaired t-tests and *p-values* are shown above each pair (ns = not significant).

We further characterized phage-resistant colonies from three representative phage-host pairs in detail: ATCC 10790 with vB_VpaP_P3, ABVat2 with vB_VatP_P5, and ABVat3 with vB_VatM_P1. Phage resistant colonies from all three pairings were sequenced using long reads and, in each case, the entire phage genome was found integrated into the host chromosome, confirming that these resistant mutants were *bona fide* lysogens (Fig. S4). The lysogens are hereafter referred to as ATCC 10790::vB_VpaP_P3, ABVat2::vB_VatP_P5, and ABVat3::vB_VatM_P1. The podophages integrate at the same chromosomal locus in their respective hosts, flanked by homologs of an undecaprenyl-diphosphatase and a KWG-repeat family protein, while the myophage integrates at a distinct locus (Fig. S4). Analysis of their *att* (attachment) sites revealed that both podophages share a core sequence (TATCGTTATCGGAAGTAAT, 19 bp) that is present in their hosts while the attachment sequence targeted by the myophage is distinct (CTGTTTATATAACATAT, 17 bp).

To assess the superinfection immunity provided by the prophages, we spotted phage on lawns of their corresponding wild type and lysogenized strains. ATCC 10790::vB_VpaP_P3 and ABVat3::vB_VatM_P1 showed complete immunity while ABVat2::vB_VatP_P5 had partial immunity with ∼30-fold greater resistance than wild type (EOP = 0.035) (Fig. 4B). Two lines of evidence suggested that phage resistance was conferred by the integrated prophage through a post-attachment mechanism, potentially involving repressor-mediated immunity, rather than altered phage adsorption. First, adsorption assays revealed no significant difference in adsorption rates between wild type and lysogens for any of the three pairs (Fig. 4C). Second, silver staining to check lipopolysaccharide (LPS) revealed no discernible difference in the staining pattern between wild type and lysogens consistent with no detectable changes to LPS (Fig. S5).

To test whether lysogeny confers a growth advantage to the host, we compared growth of ATCC 10790 and ABVat3 along with their lysogens across multiple media conditions: rich media (SK), nutrient-deplete media (10% SK), 25% saliva (v/v in water) and 25% saliva supplemented with 0.6% lactate, a condition representative of the oral environment where *Veillonella* uses lactate as its primary carbon source (42). We found no differences in growth rates between WT and their lysogens in any tested media condition (Fig. S6). Thus, the prophages either provide no apparent growth advantage to their lysogenized host, or they provide some benefit but under conditions in the natural environment that we are unable to mimic *in vitro*.

### Novel *Veillonella* prophages are spontaneously induced during exponential growth

To determine whether the *Veillonella* prophages are induced during exponential growth, lysogens were grown in rich media and supernatants were harvested at multiple time points, filtered, and spotted on lawns of the wild type host to measure phage titers over time. We found that the podophages accumulated in the supernatants of their respective lysogens over time during exponential growth (Fig. 5A). vB_VpaP_P3 reached a peak titer of ∼1.2×10⁵ PFU/ml by 2-4 hours while vB_VatP_P5 reached a peak of ∼3.7×10⁸ PFU/ml by 10 hours. Phage titers in the supernatant increased steadily during exponential growth and plateaued during stationary phase. Notably, addition of mitomycin C, a DNA damaging agent that activates the SOS response and triggers induction of many temperate phages, had no effect on production of infectious phages (Fig. S7). However, we did not establish that mitomycin C induces the SOS response in these strains. In contrast, we detected no infectious vB_VatM_P1 phage in supernatants from its lysogen during exponential growth or in stationary phase (Fig. 5A). Exposure to a range of stress conditions or different ratios of susceptible wild type cells in the environment also failed to produce detectable infectious phage release (Fig. S8A, B). Together, these results demonstrate that some *Veillonella* prophages spontaneously induce during exponential growth under nutrient rich conditions.

**Figure 5.**
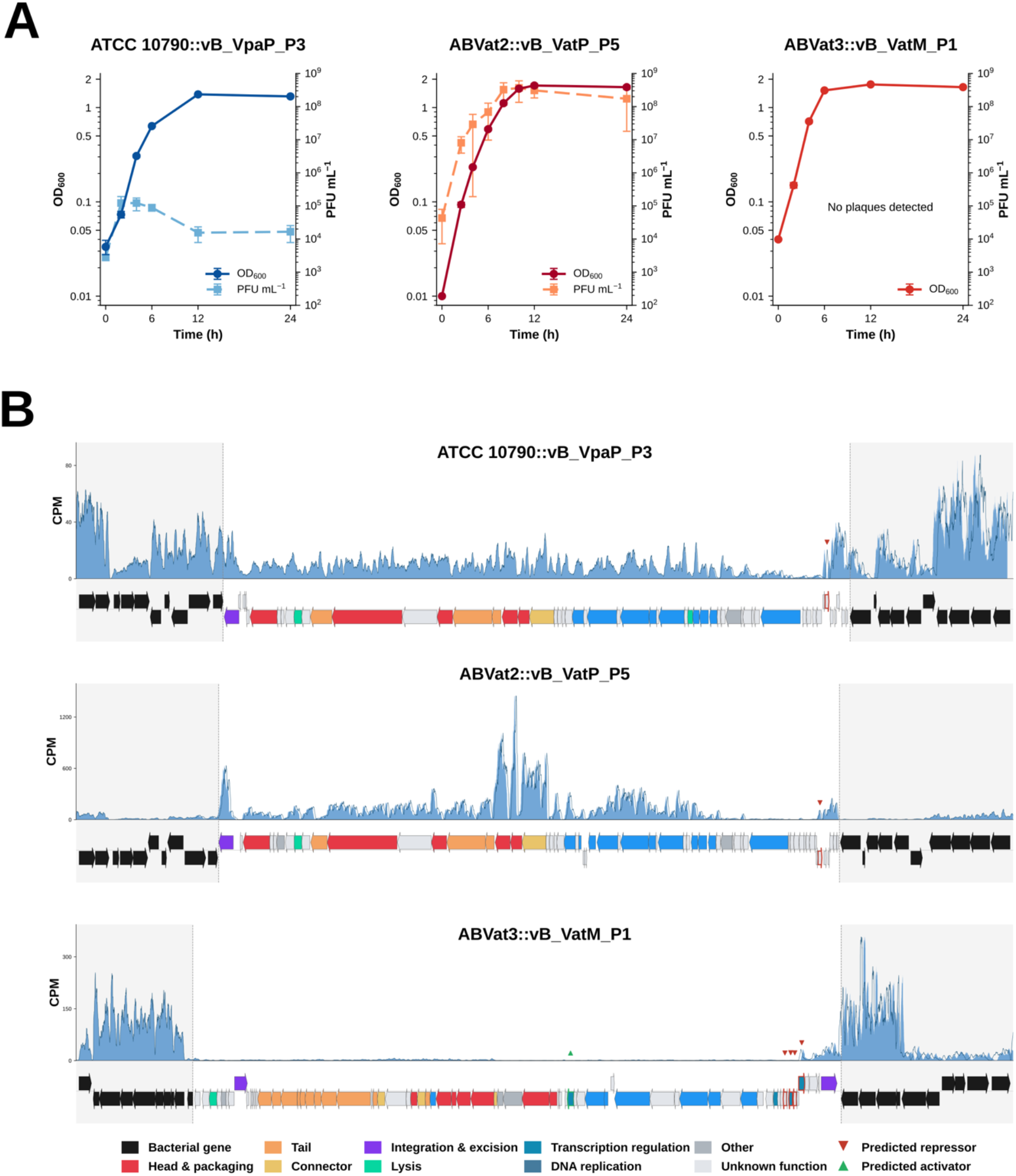
*Veillonella* lysogens spontaneously induce during exponential growth. (A) Graphs showing the free phage over time in the culture supernatants of *Veillonella* lysogens depicting spontaneous induction. Growth curves (left axis) of the cultures are shown overlaid. No PFU were detected in supernatants of ABVat3::vB_VatM_P1 at any time point. (B) RNA-seq coverage (counts per million, CPM) across the integrated prophage region and 10 flanking host genes on either side for each of the three lysogens are shown. Genes are colored by functional gene category as in Fig. 2B. Host genes are black and shaded. Repressor and activator genes, as predicted by Empathi and Pharokka are marked with triangles.

To directly assess prophage transcriptional activity in the three lysogens, we performed RNA-seq on exponentially growing cultures. We found that phage gene expression levels differed dramatically across lysogens and were strikingly consistent with each lysogen’s spontaneous induction phenotype (Fig. S9). ABVat2::vB_VatP_P5 showed the highest phage gene expression (mean 693 TPM, median 436 TPM), while ATCC 10790::vB_VpaP_P3 showed intermediate expression (mean 52 TPM, median 41 TPM) and ABVat3::vB_VatM_P1 showed the lowest (mean 15 TPM, median 8 TPM). This result indicates that transcriptional activity across the prophage is substantially elevated in lysogens with higher levels of infectious phage release. However, because these are bulk measurements, they report a population average and cannot distinguish uniform prophage expression across all cells from high-level expression confined to a subpopulation. Per-gene coverage profiles illustrated these differences in greater detail (Fig. 5B). In ABVat2::vB_VatP_P5, the counts per million (CPM) signal was broadly elevated across the phage genome with genes encoding head and connector proteins at the middle of the prophage showing highest expression. This pattern is consistent with spontaneous prophage induction but does not establish the proportion of cells undergoing induction. Interestingly, the integrase gene at the phage-host boundary was also highly expressed. In ATCC 10790::vB_VpaP_P3, a similar broad pattern was observed but at lower levels, consistent with lower overall infectious phage production in this lysogen. By contrast, in ABVat3::vB_VatM_P1 phage gene expression was largely confined to the putative repressor and integrase module, with structural and lytic genes showing minimal expression, consistent with maintenance of the lysogenic state under these conditions.

### Phage resistant mutants identify the lipopolysaccharide O antigen as the receptor for the lytic phage *vB_VnaM_P1*

In contrast to the temperate phages, whose phage resistant survivors were lysogens protected by superinfection immunity, the strictly lytic phage vB_VnaM_P1 lacks an annotated integrase and cannot establish lysogeny; hence, phage resistant colonies are likely to be receptor mutants. We exploited this distinction to identify the vB_VnaM_P1 receptor. Host strain ABVna2 was challenged with high titer vB_VnaM_P1, and five putative resistant colonies were isolated and confirmed for stable resistance by spot assay. Out of the five resistant mutants, four showed complete immunity (ABVna2_mut1-mut4) while the fifth showed only partial resistance (Fig. 6A).

**Figure 6.**
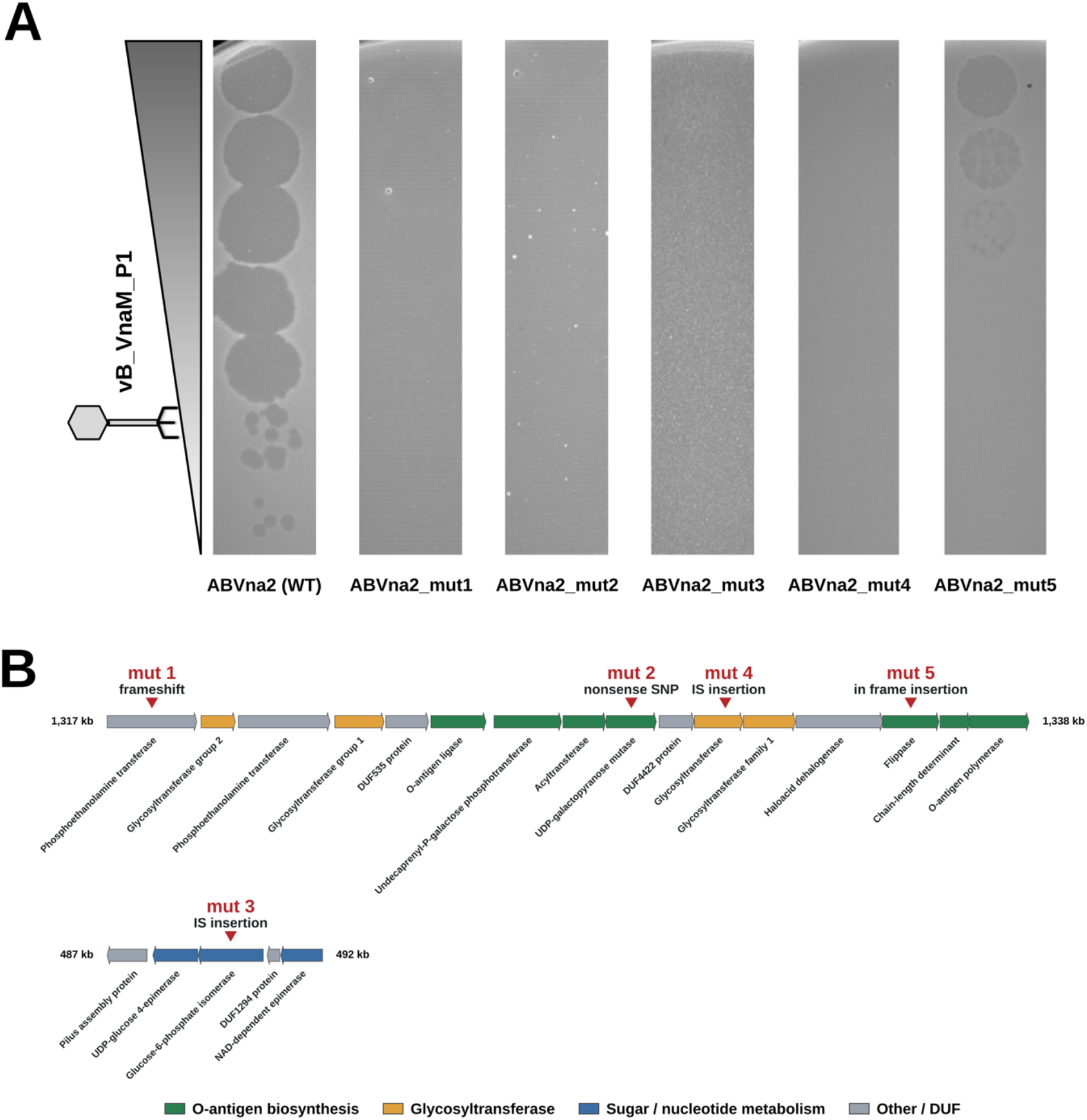
LPS O-antigen of *ABVna2* is the likely receptor of phage vB_VnaM_P1. (A) Spot assay of vB_VnaM_P1 on lawns of wild type ABVna2 and five independently isolated resistant mutants (ABVna2_mut1–5), showing reduced or abolished plaque formation in the mutants. (B) Mapping of resistance conferring mutations onto the host genomic loci implicated in vB_VnaM_P1 resistance. The O-antigen biosynthesis cluster (∼1,317–1,338 kb) and a second sugar metabolism associated locus (∼487–492 kb) are indicated and genes are colored by predicted function. The mutated gene and type of mutation indicated.

Long read sequencing revealed that each isolate carried a mutation which mapped independently to a gene required for LPS O-antigen biosynthesis (Fig. 6B). Four mutants carried lesions in the dedicated O-antigen assembly locus: an IS (Insertion Sequence) element insertion in a putative glycosyltransferase (ABVna2_mut4), an in-frame insertion in the O-antigen flippase (ABVna2_mut5), a nonsense mutation in the UDP-galactopyranose mutase (ABVna2_mut2) and a frameshift in a phosphoethanolamine transferase (ABVna2_mut1). The in-frame mutation in ABVna2_mut5 adds an additional phenylalanine residue in the flippase which might partially reduce its efficiency leading to less overall O-antigen on the surface and hence reduced but not abolished infectivity (Fig. 6A). The remaining mutant (ABVna2_mut3) carried an IS element insertion in a glucose-6-phosphate isomerase at a separate genomic locus. This gene is involved in carbohydrate metabolism and may contribute to the synthesis of nucleotide-sugar precursors required for O-antigen biosynthesis. The recovery of independent mutations in multiple dedicated O-antigen synthesis or precursor-generating genes strongly suggests that O-antigen serves as the receptor for phage vB_VnaM_P1.

### Relatives of novel phages are present at variable abundance in oral metagenomes

To check whether phages related to our isolates are present in oral metagenomes, we compared the seven phage genomes by BLASTN against three public catalogues: the Oral Phage Database (OPD 2025, 189,859 contigs), the Oral Virome Database (OVD 2022, 48,425 genomes) and the Human Oral Virome Database (HOVD, 24,440 genomes). Out of 262,724 total references, 30,912 (11.8%) produced an alignment to at least one phage but none reached both 95% nucleotide identity over 95% query coverage, indicating that no catalogue contains the same exact phage (Fig. 7A). Applying a relaxed criterion of ≥90% identity across ≥90% of the query, vB_VatM_P1 had the most related reference genomes (60 references), followed by vB_VatP_P4 (39 references), vB_VatP_P5 (34 references), vB_VpaP_P1 (23 references), vB_VatS_P6 (4 references), vB_VpaP_P3 (3 references), and vB_VnaM_P1 had none. The highest identity recorded for any reference meeting both thresholds was 94.3%, between vB_VatP_P4 and a 41.2 kb contig present in both HOVD and OVD 2022. Host predictions were available for only 82 of the 163 references passing the threshold and of these, 74 included *Veillonella* and eight *Prevotella*.

**Figure 7.**
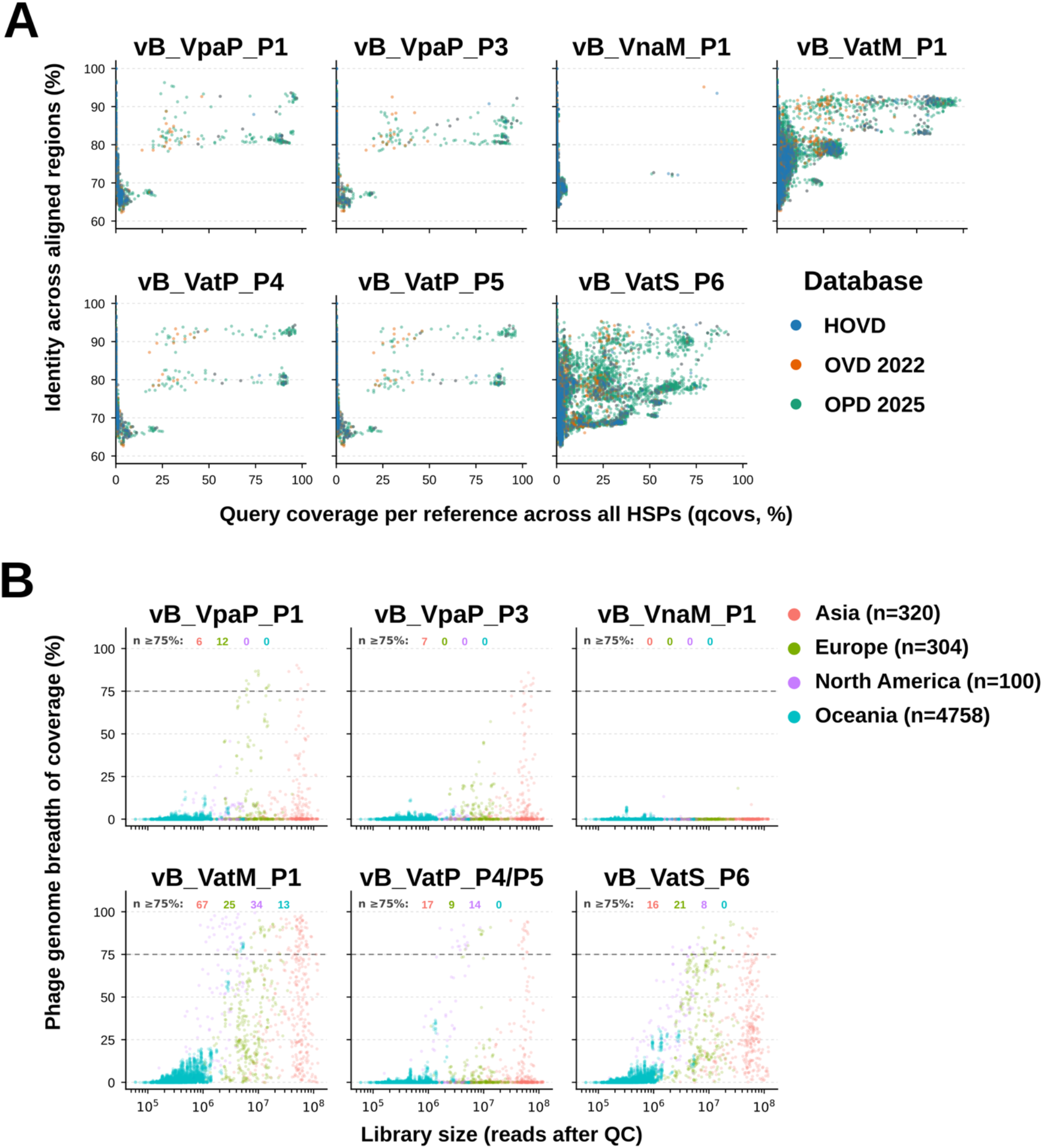
Relatives of novel *Veillonella* phages are represented in public oral virome catalogues and metagenomes. (A) Each isolated phage genome was searched by BLASTN against 262,724 reference sequences from three oral virome catalogues (HOVD, OVD 2022 and OPD 2025). Each point is one phage-reference pair; the x axis is the proportion of the phage genome covered by all HSPs (High Scoring Pairs) against that reference and the y axis is nucleotide identity computed across all aligned positions of that pair. No reference in any catalogue reaches both ≥ 95% identity and ≥ 95% coverage. (B) Reads from 5,482 publicly available oral metagenomes were mapped to the phage genomes. Each point is one metagenome, with library size after quality filtering on the x axis and the proportion of the phage genome covered at ≥ 1× on the y axis; color denotes the geographic region of the parent study and sample numbers per region are given in the legend. The dashed line marks the 75% breadth-of-coverage threshold used here to call a detection, and the counts within each panel give the number of metagenomes in each region reaching it. vB_VatP_P4 and vB_VatP_P5 are shown merged because one genome is contained within the other at 100% identity, so reads cannot be assigned between them.

Absence of a near identical catalogue genome need not mean absence from oral communities, since low abundance phages assemble in fragments and are under-represented in assembly derived catalogues (44). We therefore mapped reads from 5,482 human oral metagenomes, comprising six studies from four continents, onto the phage genomes and scored detection as breadth of coverage: the proportion of each genome covered by at least one read. At least one read mapped in 11.5-80.8% of samples, but the median breadth among these was only 0.13-1.6%. Requiring ≥75% of the genome to be covered at ≥1× depth (44), vB_VatM_P1 was the most frequently detected phage and was detected in 139 samples, followed by vB_VatS_P6 (45 samples), vB_VatP_P4/P5 (40 samples), vB_VpaP_P1 (18 samples), vB_VpaP_P3 (7 samples) and vB_VnaM_P1 was detected in none (Fig. 7B). This ranking of detection frequency was unchanged at breadth thresholds of 25% and 50%. At the defined threshold, detections were well supported, with a median genome coverage of 5.3× (range of 1.7× - 182.6×). Where detected, however, the phages were minor components of the metagenome with a median of 161 reads per million and no phage accounted for more than 0.22% of the reads in any library. Geographically, detections spanned all four continents for vB_VatM_P1 while vB_VpaP_P3 was only detected in the Asian samples. Comparisons between regions are confounded, however, since sequencing depth differed 160-fold between studies. Nonetheless, we noticed a location specific difference between the phages vB_VatM_P1 and vB_VatS_P6: in the Oceanian samples exceeding 3 million reads, vB_VatM_P1 reached a median breadth of 79.5% while vB_VatS_P6 reached only 11.0%, whereas the two phages had comparable abundances in Asian (39.8% and 25.7%) and European (26.3% and 35.7%) samples of similar depth. Together these results indicate that the isolated phages are novel and not represented in current oral virus catalogues, yet related sequences are recoverable from oral metagenomes in geographically distinct locations.

## Discussion

This study presents the first systematic isolation and genomic characterization of phages infecting *Veillonella* which play an important role in the human oral microbiome. Despite the relevance of *Veillonella* as an emerging accessory pathogen, no phages infecting this genus had previously been characterized at the genomic and functional level. Here we describe seven unique, novel phages infecting natural *Veillonella* isolates from three different species and reveal diversity in genome organization, host range, and lifestyle. All except one of the isolated phages have an annotated integrase gene and appear to have a temperate lifestyle consistent with reports that most phages identified in the oral cavity are predicted to be temperate (45, 46). Among the isolated phages, the podophages infecting *V. parvula* or *V. atypica* isolates fell into a distinct cluster and may represent one or more novel genera in the *Autographiviridae* family. The myophages and siphophage were distantly related to any phage in the Virus-Host DB and are potentially the first identified members of their own novel genera.

Genomic characterization revealed the phages harbor a diverse repertoire of genes that reflect specialization to their host and lifestyle. All the podophages encode their own type III anaerobic ribonucleotide reductase which might support DNA replication in anaerobic subgingival plaque, a niche preferred by *V. parvula* (47). The myophage vB_VatM_P1 encodes DNA methyltransferase subunits that could help it evade host RM systems including the type I, II and IV systems present in ABVat3. The putative lytic phage vB_VnaM_P1 encodes its own nucleotide metabolism toolkit, including two DNA primases, a dUTPase, a nucleotide kinase, and a thymidylate synthase, consistent with the strategy among lytic phages of supplementing host dNTP pools to sustain rapid genome replication (48, 49). It also encodes an Erf like ssDNA annealing protein, suggesting it may use recombination mediated replication, another strategy common among fast replicating lytic phages (49). Strikingly, the siphophage *vB_VatS_P6* encodes a module suited for autonomous, extrachromosomal replication including a replication initiation protein, a DnaC like helicase loader, a RecT like ssDNA annealing protein, and a RusA like Holliday junction resolvase. We also identified a Leu tRNA targeting attachment site in the phage, suggesting it can integrate into its host genome as well. Further work is needed to elucidate whether this phage is maintained as a chromosomally integrated prophage or as an extrachromosomal episome and what are the physiological consequences for its host.

We showed that lysogens formed readily *in vitro* for all three natural phage-host pairs tested, and in each case the lysogen resisted superinfection after the phage had adsorbed, consistent with repressor mediated immunity. Nonetheless, the three prophages behaved differently with regard to induction which may be linked to the architecture of their lysis-lysogeny switches (Fig. S10 and Table S7, S8). vB_VatM_P1, which released no detectable infectious phage under any condition tested, encodes a CI-type repressor transcribed divergently from a DUF739 family protein homologous to MOR, the antirepressor of Lactococcus phage TP901-1 and *S. aureus* phage φ13 (50, 51). In that switch, when the prophage commits to lysis, MOR binds the N-terminal domain of CI and blocks it from occupying its operator sites and the resulting CI:MOR complex is thought to repress the lysogenic promoter (51, 52). In the ABVat3 lysogen, the antirepressor gene was barely transcribed while the repressor was expressed seventeen-fold above the prophage median (Fig. 5B), consistent with a predominantly repressed lytic program. Whether this predicted antirepressor controls induction of vB_VatM_P1, and what signals might activate it, remain unknown. The siphophage vB_VatS_P6 shares a similar architecture but different components: the gene transcribed divergently from its CI-type repressor encodes a second predicted repressor rather than a MOR homologue, and an antirepressor of the unrelated ANT family lies 6.8 kb away within the replication module (Fig. S10). The lysogenic podophages encode a simpler module consisting only of a single small repressor (Fig. S10). We showed that podophage lysogens induced spontaneously, but to very different extents (Fig. 5A). This parallels their superinfection immunity, which was complete for ATCC 10790::vB_VpaP_P3 and only partial for ABVat2::vB_VatP_P5. Differences in repressor activity could contribute to these phenotypes, although the underlying regulatory mechanisms have not been established. Consistent with this, the two repressors share only 57% amino acid identity. Why these different induction strategies exist and the advantage each provides in the context of the oral biofilm remain to be determined.

Our results place these phages where oral virome sequence data and phage biology meet. Sequences related to the isolates are recoverable from oral metagenomes on several continents, yet none of the three oral virus catalogues contains a close match to our isolated phages, and many related sequences lack confident host assignments. Bringing them into culture makes accessible the properties that sequence alone cannot report: host range, integration site, superinfection immunity, spontaneous induction and receptor identity. Because *Veillonella* participate in extensive metabolic and physical interactions within oral biofilms, phages that shape *Veillonella* populations may also influence the structure of the wider community.

## Materials and methods

### Study participants and IRB

Oral rinse samples were collected under a protocol approved by the University of Minnesota Institutional Review Board (STUDY00025333; principal investigator, Julia Willett). The study, titled “D2D Collection of microbes from the oral cavity using the oral rinse method,” was approved on May 22, 2025, with an effective date of June 18, 2025. Samples were collected with informed consent from participants.

### Strains and growth conditions

All strains, phages and primers used in this study are listed in Supplementary Tables. *Veillonella* strains were routinely grown in SK (54) or BHIL (13) medium at 37 °C without shaking in a Coy anaerobic chamber (80% N₂, 10% H₂ and 10% CO₂), unless otherwise indicated. Oral *Veillonella* Selective Media (OVSM) (55) was used for isolation from clinical samples.

### Isolation of novel *Veillonella* strains

Pooled oral rinse samples were centrifuged (5,000 rcf, 10 min) and the cell pellet was streaked for single colonies on OVSM. Colonies were restreaked, grown overnight in SK, and boiled lysates were screened by PCR with *Veillonella* specific *rpoB* primers, with universal 16S primers as a lysis control (Table S9). Full detail is given in Supplemental Methods.

### Isolation of novel *Veillonella* phages

Oral rinse samples in PBS were clarified and filtered (0.45 µm). Filtrate (1–2 ml) was enriched for 24 h in 9 ml SK containing 100 µl overnight host culture and 1 mM each CaCl₂ and MgCl₂, then clarified and filtered as above. Enrichments were screened for plaques by double agar overlay, and individual plaques were picked into TSM buffer (40 mM Tris, 68 mM NaCl, 10 mM MgCl₂) and purified through at least three rounds of plaque isolation before propagation in SK (Supplemental Methods).

### Phage spot assay

Lawns were prepared by mixing 100 µl overnight culture and 1 mM each CaCl₂ and MgCl₂ with molten 0.6% SK top agar and pouring over a 1.5% SK agar base. Ten-fold serial dilutions of phage (2–3 µl) were spotted, dried and incubated overnight at 37 °C. For host range and efficiency of plaquing (EOP), the titer on each test strain was taken from the highest dilution giving countable plaques and normalised to the titer on the isolation host.

### Bacterial growth assays

Overnight cultures were diluted 1:50 into the indicated medium (SK, 10% SK, 25% saliva, or 25% saliva with 0.6% lactate) in 96-well plates (Celltreat #229597) and grown anaerobically at 37 °C without shaking, with OD₆₀₀ read every 20 min for 24 h in a BioTek plate reader. Growth rate (μ) was taken as the maximal slope of ln(OD₆₀₀) over a sliding six-point window, doubling time as ln(2)/μ, and lag time by the tangent-line method at the point of maximal growth rate.

### Autoaggregation assay

Overnight cultures were normalised to OD₆₀₀ = 1 and resuspended in 1 ml aggregation buffer (1 mM Tris-HCl pH 8.0, 0.1 mM CaCl₂, 0.1 mM MgCl₂, 150 mM NaCl). Cells settled statically in a 1-cm cuvette at room temperature, and OD₆₀₀ was read from the upper portion of the suspension at time 0 and at intervals over 24–48 h. Autoaggregation is expressed as the percentage of initial OD₆₀₀ remaining in suspension.

### Isolation of resistant mutants

Phage-resistant mutants were selected either in a 0.6% SK agar overlay seeded with high-titer lysate, or by challenging an exponential-phase culture with lysate overnight and plating dilutions on SK. Surviving colonies were restreaked and confirmed resistant by spot assay before sequencing.

### Adsorption assays

Overnight host cultures were mixed with phage at an MOI of 0.05–0.1 in 1 ml SK containing 1 mM each CaCl₂ and MgCl₂; SK without cells served as the no cell control. After 20 min at 37 °C, cells and adsorbed phage were removed by centrifugation and filtration (0.22 µm), and unadsorbed phage in the filtrate was titered by plaque assay. Percentage adsorption is expressed relative to the no cell control.

### Induction assays

For spontaneous induction, overnight lysogen cultures were diluted 1:50 in SK and incubated at 37 °C; 1 ml aliquots taken at intervals were clarified (15,000 rcf, 2 min), filtered (0.22 µm) and spotted on lawns of the wild-type host. For chemical induction, exponential-phase cultures were treated with mitomycin C (4 µg/ml) or an equal volume of DMSO vehicle and titers were followed over time. vB_VatM_P1 was additionally tested against a panel of eleven stress conditions and three lysogen:wild type coculture ratios, with supernatants assayed both by spot test and by PCR for the phage integrase gene (Table S9). Conditions, incubation times and controls are given in Supplemental Methods.

### Efficiency of lysogeny

Ten-fold serial dilutions of overnight host cultures were spread on SK plates seeded with or without 100 µl high-titer phage lysate. After three days at 37 °C, the fraction of colonies surviving on phage-seeded plates relative to unseeded control plates was used as an estimate of efficiency of lysogeny.

### Genomic DNA sequencing, assembly and annotation

Bacterial genomic DNA was extracted with the Qiagen DNeasy kit using bead-beating lysis. Phage DNA was prepared from high-titer lysates by nuclease treatment, PEG precipitation and proteinase K digestion, followed by extraction with the Promega Wizard kit (Supplemental Methods). Genomes were sequenced by Oxford Nanopore (Plasmidsaurus), PacBio HiFi (UIUC Sequencing Core) or Illumina (SeqCenter). Long reads were filtered with Filtlong v0.3.1 (lowest-quality 5% of reads and reads <1,000 bp removed) and assembled with Autocycler v0.5.2 (56). Bacterial assemblies were annotated with Bakta v1.11.4 (57). Illumina reads were filtered with fastp v1.1.0 (58) and assembled with Unicycler v0.5.1 (59) using default settings. Reads for vB_VnaM_P1 were downsampled to 1,000× coverage before assembly with SPAdes v4.3.0 (60). Phage genomes were annotated with Pharokka v1.9.1 (61) and Empathi v1.0.6 (62). Anti-phage defence systems were identified with PADLOC (63).

### RNA sequencing and analysis

Lysogens were diluted 1:100 into 10 ml SK, grown to exponential phase, washed twice in PBS to remove free phage and lysed by bead beating. Total RNA was extracted with the Qiagen RNeasy kit, DNase-treated, and sequenced by SeqCenter on a NovaSeq X Plus (Illumina Stranded Total RNA Prep with Ligation, Ribo-Zero Plus; 150 bp paired-end). Reads were trimmed with fastp v1.1.0 (58) and aligned to the corresponding lysogen genome with Bowtie2 v2.5.5 (64). Alignments were coordinate-sorted and indexed with SAMtools v1.23.1 (65). CPM-normalised coverage tracks (5 bp bins) were generated with bamCoverage from deepTools v3.5.6 (66). Read pairs over CDS features were counted with featureCounts v2.1.1 (67) in strand-specific mode, using Bakta GFF3 annotations supplemented with the coordinates of Pharokka-annotated phage CDS features absent from the Bakta annotation. Transcript abundance is reported as transcripts per million (TPM).

### Comparative genomics and Phylogenetic classification

Bacterial assemblies were placed in the GTDB reference tree using GTDB-Tk (v2.6.1, release 226) (68) to determine their relationship to reference *Veillonella* strains. The tree was rooted on *V. agrestimuris* and *V. caviae,* the deepest branching lineage. Average nucleotide identity between bacterial genomes and, separately, between phage genomes was calculated with FastANI (v1.3.4) (69). Phage genomes were compared against the Virus-Host DB with ViPTree (70), which scores pairwise similarity as Sg (36), a normalised tBLASTx score ranging from 0 to 1. Synteny and pairwise nucleotide identity among Pharokka-annotated phage genomes were visualised with clinker (71), with genes coloured by Pharokka functional category (Fig. 2B).

### Evaluating homology between isolates and known phage

Seven isolated phage genomes were compared against the Human Oral Virome Database [10.1016/j.xcrm.2025.102325], Oral Virus Database [10.1016/j.isci.2022.104418], and Oral Phage Database [10.1038/s41522-025-00773-z] reference collections using NCBI BLAST+ v2.15.0+. Nucleotide databases were constructed using makeblastdb with -dbtype nucl - parse_seqids -blastdb_version 5. Searches were performed using blastn with -task blastn - word_size 11 -evalue 10 -dust yes -soft_masking true. For each subject, a genome-wide identity was calculated as the mean percent identity weighted by high-scoring segment pairs alignment length. Subjects were ranked by the product of coverage and length-weighted identity and the ten highest-ranked retained for each phage.

### Quantifying abundance of phage in publicly available oral metagenomes

Raw oral metagenomic reads (ERP006678, ERP110622, ERP169403, SRP029441, SRP072561, and SRP133047) were downloaded from the sequence read archive and cleaned using <u>fastp</u> v1.3.1 [10.1093/bioinformatics/bty560] with default parameter --qualified_quality_phred 15, -- unqualified_percent_limit 40, --n_base_limit 5, --length_required 15, --average_qual 0. Cleaned reads were mapped separately against a joint index of the seven isolated phage genomes using <u>BBSplit</u> from BBTools v39.38. Mapping used default alignment-sensitivity settings, and qtrim=false, ambiguous=best, ambiguous2=best, secondary=false, secondarycov=false, physcov=false, delcoverage=false, 32bit=true, and nzo=false. Paired reads mapping to each phage were normalized using total sum scaling. For each phage, differences between geographic region were quantified using a Kruskal-Wallis test and a Pairwise Wilcoxon Rank Sum compared between group levels (holm correction) in R (v4.6.0). Similarly, presence/absence across samples was modeled as a function of region using logistic regression (generalized linear model with a binomial error distribution and logit link) in R. A separate model was fitted for each phage, and the overall effect of region was evaluated by analysis of deviance, comparing each fitted model to the corresponding intercept-only model by likelihood-ratio test (χ², [N−1] df).

### Phage genome packaging and termini prediction

Virion reads were mapped to each assembly with minimap2 (v2.31) (72) (‘-ax map-hifì for PacBio HiFi, ‘-ax sr’ for Illumina) and sorted with SAMtools (v1.24) (65); duplicates were removed from the Illumina libraries with ‘samtools markdup -r’. Per-base depth from primary alignments was normalised to the contig median, and the 5′-most aligned reference base of each primary alignment was counted separately per strand, excluding the five terminal bases of each contig. A direct terminal repeat was called where an interval carried approximately twice the flanking depth and was bounded at both edges by single-base, strand-specific read-end pileups.

### Identification of lysis-lysogeny switch genes

Candidate transcriptional regulators among Pharokka-annotated coding sequences were predicted with Empathi (62) at a score threshold of ≥ 0.95, with repressor and activator roles read from the corresponding channels (Tables S7 and S8). Because that prediction is not homology-based, candidates were searched for conserved domains with hmmscan (73) against Pfam-A (74). Where a domain assigned a protein to a characterized phage regulator family, the corresponding reference protein was retrieved from GenBank by accession and the relationship confirmed by pairwise alignment with phmmer and blastp. Proteins were assigned to a family only where a Pfam domain was detected and candidates supported by the Empathi prediction alone were reported without a family assignment. Integrase genes and attachment sites were taken from the Pharokka annotation.

### Lipopolysaccharide (LPS) silver staining

Whole-cell lysates of plate-grown *Veillonella* were digested with proteinase K, resolved on a 15% SDS-PAGE gel and visualized by periodate oxidation followed by ammoniacal silver staining (Supplemental Methods).

### Identification of resistance-conferring mutations

Genomic DNA from phage-resistant mutants was sequenced and assembled as described above. Mutant genome assemblies were aligned to the corresponding wild type reference genome using Mauve (75) to identify mutations.

### Transmission electron microscopy (TEM)

Phage lysates were concentrated, washed in 0.1 M ammonium acetate and DNase-treated (Supplemental Methods). Samples were negatively stained with 2% uranyl acetate on carbon-coated grids and imaged at 120 kV on a JEOL JEM-1400 (vB_VpaP_P1, vB_VpaP_P3, vB_VatM_P1, vB_VatP_P4 and vB_VatP_P5) or on an FEI Tecnai Spirit (vB_VatS_P6 and vB_VnaM_P1). Capsid diameters (the mean of two diagonal measurements per particle) and tail lengths were measured in Fiji (76) for at least 10 particles per phage (n = 5 for vB_VatS_P6) and are reported as mean ± standard deviation.

### Figures and statistics

All figures were generated in Python 3.13 with Matplotlib and exported as vector graphics. Adsorption data were compared by two-tailed unpaired *t* test, with p values shown in Fig. 4C. Central tendency is reported as median with interquartile range for distributions that are not symmetric and as mean ± standard deviation otherwise. The sample number (n) is stated in each figure legend and refers to biological replicates throughout.

## Acknowledgements

We thank Julia Willett and members of the Willett laboratory at the University of Minnesota for their collaboration in collecting oral rinse samples. We are grateful to Mark Herzberg and Karen Johnstone at the University of Minnesota School of Dentistry for valuable discussions and insights into oral microbiology and oral microbial communities. We also thank Asma Hatoum-Aslan, Michael Miller, and members of their laboratories at the University of Illinois Urbana-Champaign for helpful discussions and scientific input, as well as members of the Vanderpool laboratory for discussions, feedback, and support throughout this work.

This research was supported by the National Institutes of Health, National Institute of General Medical Sciences (R35GM139557 to C.K.V.), and institutional funds from the University of Minnesota.

Claude (Anthropic) was used as a computational assistant during data analysis and manuscript preparation. Its use covered writing and debugging analysis and plotting code, reproducing and independently verifying analyses performed, tabulating annotation and alignment output, and editing text for concision and clarity. All experiments were designed and performed by the authors. All analyses were specified by the authors, and every quantitative result reported here was derived from the underlying data and checked against the source files by the authors, who take full responsibility for the accuracy and integrity of the work.

## Conflict of interest

The authors declare no conflict of interest.

## Supplementary material

Supplementary tables S1-S9.

Supplementary figures S1-S10.

## Supplementary Methods

## Data availability

All raw sequencing data and assembled genomes in this study are available in NCBI under PRJXXXXX.

## Notes

### Competing Interest Statement

The authors have declared no competing interest.

