## Supplementary methods and figures for "Isolation and characterization of diverse *Veillonella* phages reveal novel lineages and distinct host interactions"

### Collection and isolation of *Veillonella* strains

Oral rinse samples were centrifuged at 5,000 rcf for 10 min to pellet cells. The pellet was resuspended in SK medium and 20 µl was quadrant-streaked for single colonies on OVSM. After two days of anaerobic incubation at 37 °C, prospective colonies were picked and restreaked on OVSM. Single colonies were then picked and grown overnight in SK. For colony PCR, 10 µl of overnight culture was mixed with 90 µl water and boiled at 95 °C for 20 min to lyse the cells, and 2 µl of this lysate was used as template for PCR with *Veillonella*-specific *rpoB* primers. Universal 16S rRNA primers were run in parallel as a lysis control. All primer sequences are given in Table S9.

### Enrichment, purification and propagation of phages

Oral rinse samples in PBS were centrifuged at 5,000 rcf for 10 min, and supernatants were filtered through a 0.45 µm filter to remove bacterial cells. For enrichment, 1–2 ml of filtrate was combined with 9 ml SK medium, 100 µl of overnight host culture and a salt solution giving 1 mM final concentration each of $CaCl_2$ and $MgCl_2$. Enrichments were incubated for 24 h, then centrifuged at 5,000 rcf for 10 min and the supernatant passed through a 0.45 µm filter.

Phage presence was tested by double agar overlay: 100 µl of each supernatant dilution was mixed with 100 µl of overnight host culture in 3 ml of molten 0.6% SK agar and poured onto a 1.5% SK agar base. After the overlay solidified, plates were incubated overnight at 37 °C and inspected for plaques. Individual plaques were picked into TSM buffer (40 mM Tris, 68 mM NaCl, 10 mM $MgCl_2$) and carried through at least three successive rounds of plaque isolation to obtain a pure preparation. After the final round, a single plaque was resuspended in 100 µl TSM and added to 10 ml SK containing 500 µl of overnight host culture and salt solution, incubated for 24 h, and then centrifuged and filtered as above. Titers of the resulting lysate were determined by serial dilution and spot assay on a lawn of the isolation host.

### Stress and coculture panel for induction of vB_VatM_P1

An overnight culture of ABVat3::vB_VatM_P1 was diluted 1:50 into SK and grown to exponential phase. The culture was divided into 3 ml aliquots and exposed to a panel of eleven conditions: mitomycin C at 0.5, 2 or 5 µg/ml; 1 mM $H_2O_2$; 1 h of oxygen exposure (37 °C, shaking); 1 h heat shock at 42 °C; 1 h cold shock at 0 °C; 15 J/m² UV; pH 4; 10% SK (starvation); and SK supplemented with 10% saliva (oral niche). To test induction in the presence of susceptible cells, overnight cultures of the lysogen were mixed with wild-type ABVat3 at ratios of 1:9, 1:1 and 9:1 (v/v).

Following treatment, aliquots were incubated for 4 h or 24 h, and supernatants were filtered and spotted onto a lawn of the wild-type host; a high-titer vB_VatM_P1 lysate served as a positive control. Supernatants were additionally screened for released phage by PCR with primers specific for the phage integrase gene (Table S9). To release phage DNA, 100 µl of supernatant was treated with 5 µl proteinase K, incubated at 56 °C for 1 h and heat-inactivated at 95 °C for 20 min, and 2 µl of the treated supernatant was used as PCR template. Phage genomic DNA and a high-titer vB_VatM_P1 lysate processed in the same way served as positive controls.

### Nucleic acid extraction

**Bacterial genomic DNA.** Between 5 and 10 ml of overnight culture was pelleted at 5,000 rcf for 10 min and the supernatant discarded. DNA was isolated with the Qiagen DNeasy kit according to the manufacturer’s instructions, with cell lysis performed by bead beating (1 mm beads, 4,000 rpm, 30 s, 2–3 rounds).

**Phage genomic DNA.** Approximately 10 ml of high-titer lysate was treated with 10 µl DNase I (NEB) and 10 µl RNase A (NEB) for 1 h at 37 °C. Phage particles were then precipitated overnight at 4 °C in PEG solution (10% PEG 8000, 1 M NaCl final concentration) and pelleted by centrifugation at 7,200 rcf for 1 h at 4 °C. The supernatant was discarded and the pellet dissolved in 500 µl of 5 mM $MgSO_4$, then treated with 5 µl proteinase K (Promega) and 5 mM EDTA for 1 h at 56 °C. DNA was extracted with the Promega Wizard DNA kit according to the manufacturer's instructions.

**Total RNA.** Overnight cultures of lysogens were diluted 1:100 into 10 ml of fresh SK and grown to exponential phase. Cells were washed twice in PBS to remove free phage and lysed by bead beating as above. Total RNA was extracted with the Qiagen RNeasy kit according to the manufacturer's instructions and treated with Invitrogen DNase I (RNase-free) to remove residual genomic DNA. Purified RNA was submitted to SeqCenter, where libraries were prepared with the Illumina Stranded Total RNA Prep with Ligation and Ribo-Zero Plus kit using 10 bp unique dual indices and sequenced on a NovaSeq X Plus to generate 150 bp paired-end reads. Demultiplexing, quality control and adapter trimming were performed with BCL Convert (v4.2.4).

### Lipopolysaccharide silver staining

*Veillonella* colonies were scraped from SK plates and suspended in ice-cold PBS to an $OD_{600}$ of 1. One millilitre of suspension was centrifuged (5,000 rcf, 5 min, 4 °C) and the pellet resuspended in 100 µl of lysis buffer (2% [w/v] SDS, 4% [v/v] β-mercaptoethanol, 10% [v/v] glycerol, 1 M Tris-HCl pH 6.8, bromophenol blue) and boiled for 10 min. After cooling to room temperature, samples were digested with 5 µl of proteinase K (20 mg/ml) overnight at 55 °C. Ten microlitres of each sample were resolved on a 15% SDS-PAGE gel.

All staining steps were performed at room temperature with shaking. The gel was oxidised for 20 min in 1% (w/v) sodium periodate, 40% (v/v) ethanol and 5% (v/v) acetic acid, then washed three times for 7 min each in $ddH_2O$. It was stained for 10 min in the dark in freshly prepared ammoniacal silver solution (0.67% [w/v] $AgNO_3$, 1.33% [v/v] ammonium hydroxide [29%], 18.7 mM NaOH), washed three times in $ddH_2O$, and developed in 0.0045% (w/v) citric acid and 0.05% (v/v) formaldehyde (37%). Development was stopped by transferring the gel to $ddH_2O$, and gels were imaged on a digital scanner.

### Preparation of samples for transmission electron microscopy

One millilitre of high-titer phage lysate was centrifuged at 21,000 rcf for 1 h at 4 °C. The supernatant was carefully discarded, leaving approximately 100 µl, and the pellet was washed with 900 µl of 0.1 M ammonium acetate. The wash was repeated twice more, and after the final spin the pellet was resuspended in the remaining 100 µl of ammonium acetate. A 17 µl aliquot of the concentrated preparation was treated with 1 µl DNase I (NEB) and 2 µl 10× DNase I buffer at 37 °C for 1 h.

For phages vB_VpaP_P1, vB_VpaP_P3, vB_VatM_P1, vB_VatP_P4 and vB_VatP_P5, a drop of the processed sample was deposited onto a glow-discharged, carbon-coated grid and incubated for 5 min. Excess liquid was removed, a drop of 2% aqueous uranyl acetate was applied for 30 s, the stain was wicked away with filter paper, and the grid was air-dried before imaging. Samples were imaged at 120 kV with an AMT NanoSprint 12L-B CMOS sensor mounted on a JEOL JEM-1400 transmission electron microscope. For phages vB_VatS_P6 and vB_VnaM_P1, samples were negatively stained with 2% uranyl acetate on 400-mesh carbon-coated grids and examined on an FEI Tecnai Spirit transmission electron microscope.

### Additional detail on packaging and termini prediction

Duplicate removal with samtools markdup -r discarded 63–65% of reads in the Illumina libraries. Per-base depth was computed from primary alignments only (samtools depth -a -Q 1 -G 0x900).

### Supplementary Figures and Figure legends

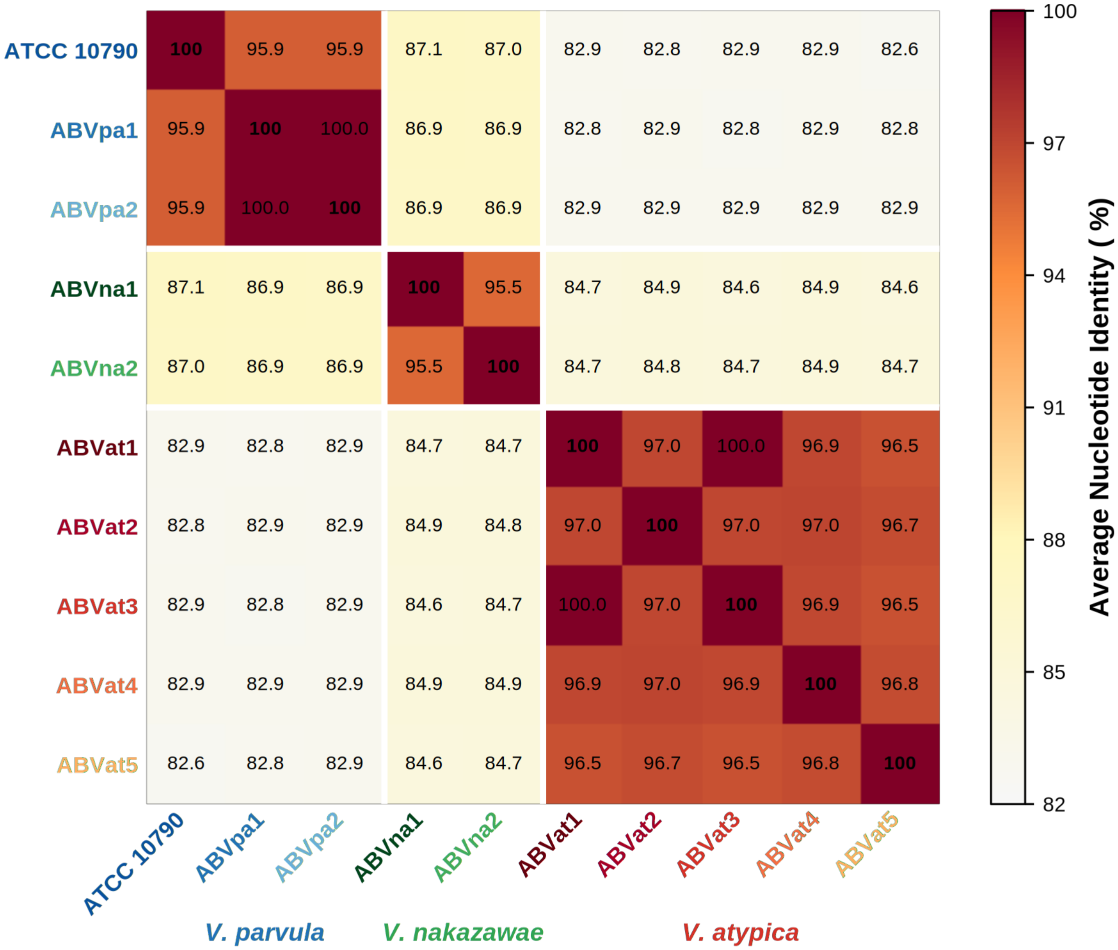


**Figure S1. Average nucleotide identity (ANI) among *Veillonella* isolates.**
Heatmap of pairwise ANI values across ten *Veillonella* strains used in this study, grouped by species. Actual values are displayed inside the heatmap tiles.

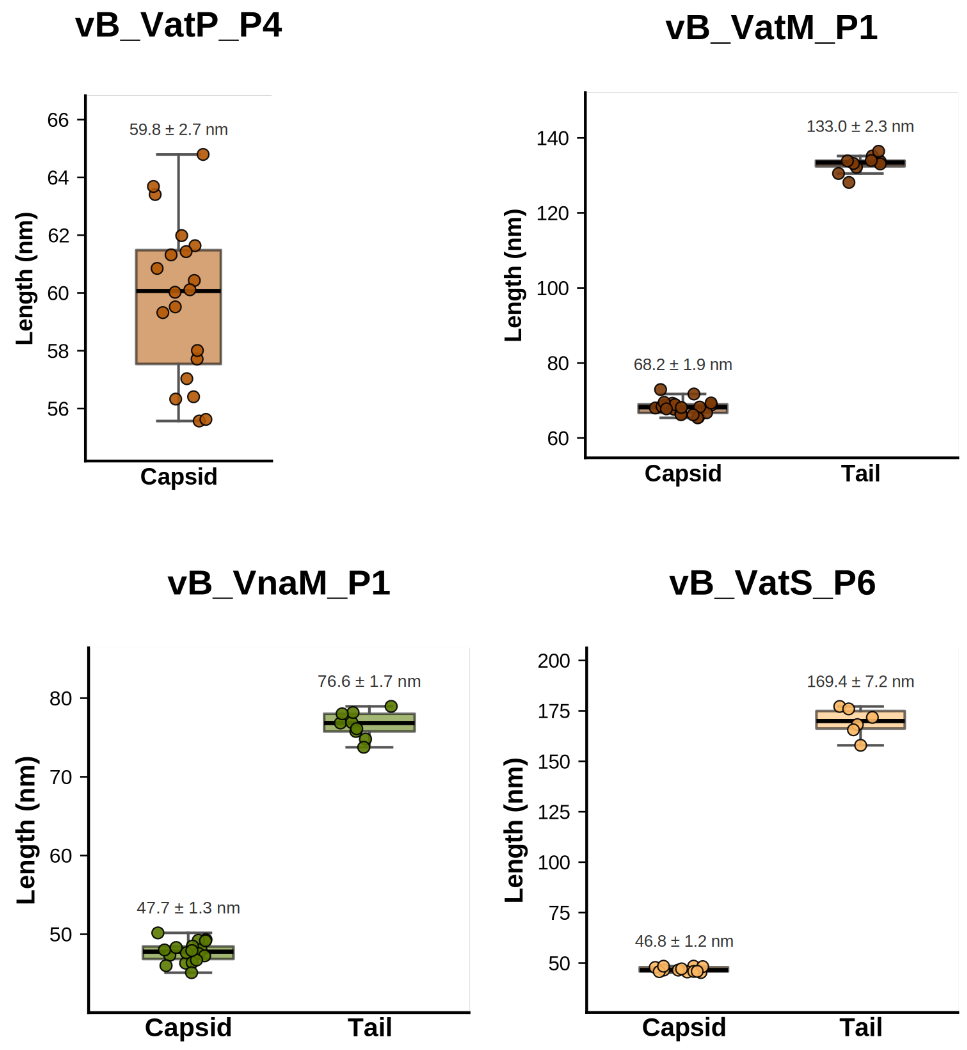


**Figure S2. Quantification of phage capsid and tail dimensions by TEM.**
Phage capsid and tail dimensions measured from transmission electron micrographs using n=10 individual phage particles (except vB_VatS_P6 tail measurements for which n=6 was used). Values are represented as a boxplot with individual data points overlaid and mean ± SD highlighted.

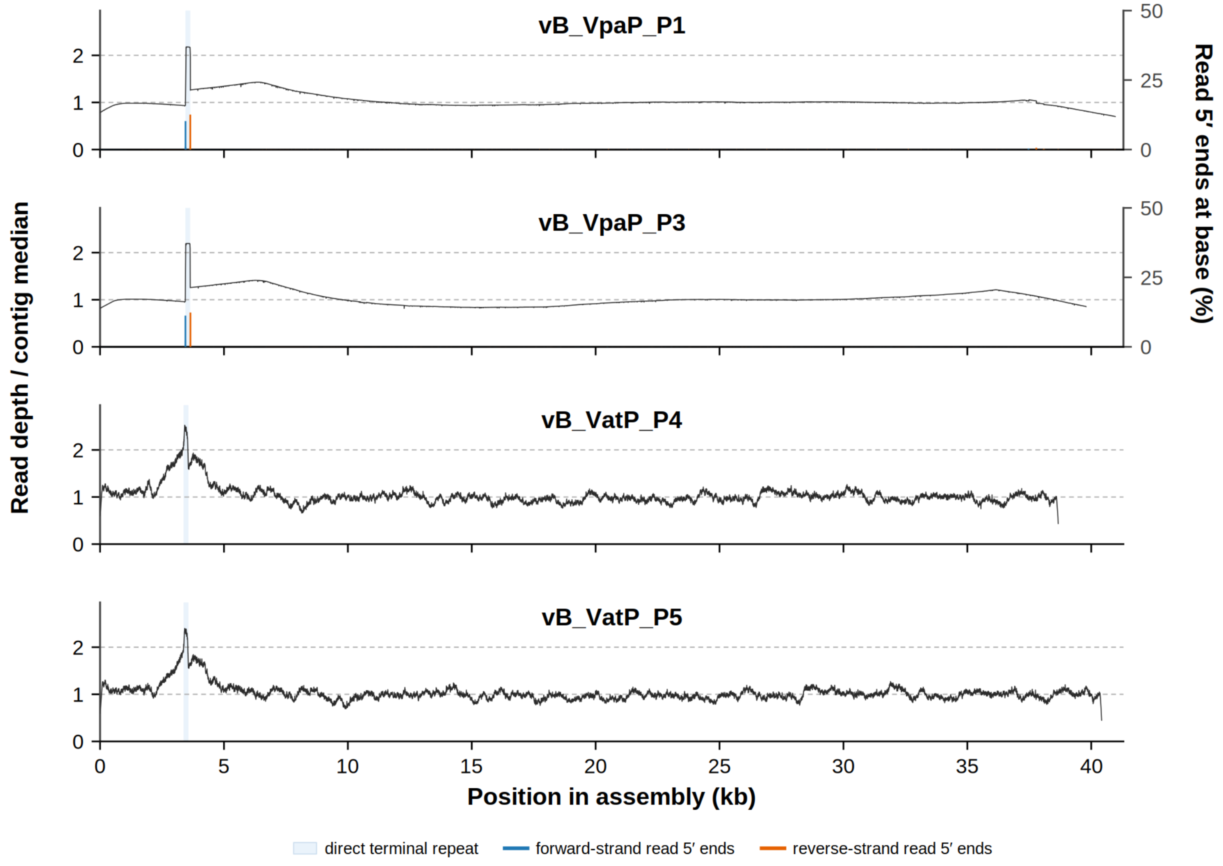


**Figure S3. Direct terminal repeats in the *Veillonella* podophages.**

Graphs showing per-base depth relative to the contig median (left axis) for the isolated podophages. In vB_VpaP_P1 and vB_VpaP_P3, sequenced with PacBio HiFi reads, the shaded 195 bp and 194 bp interval carries approximately twice the flanking depth and is bounded at both edges by single-base pileups of strand-specific read 5′ ends (right axis, % of all read ends on that strand), identifying it as a repeat present once at each end of a linear chromosome. The homologous interval in vB_VatP_P4 and vB_VatP_P5 is the highest-coverage position in each genome at a similar depth ratio, but random fragmentation during short-read library preparation leaves no read-end signal, so its boundaries are unresolved.

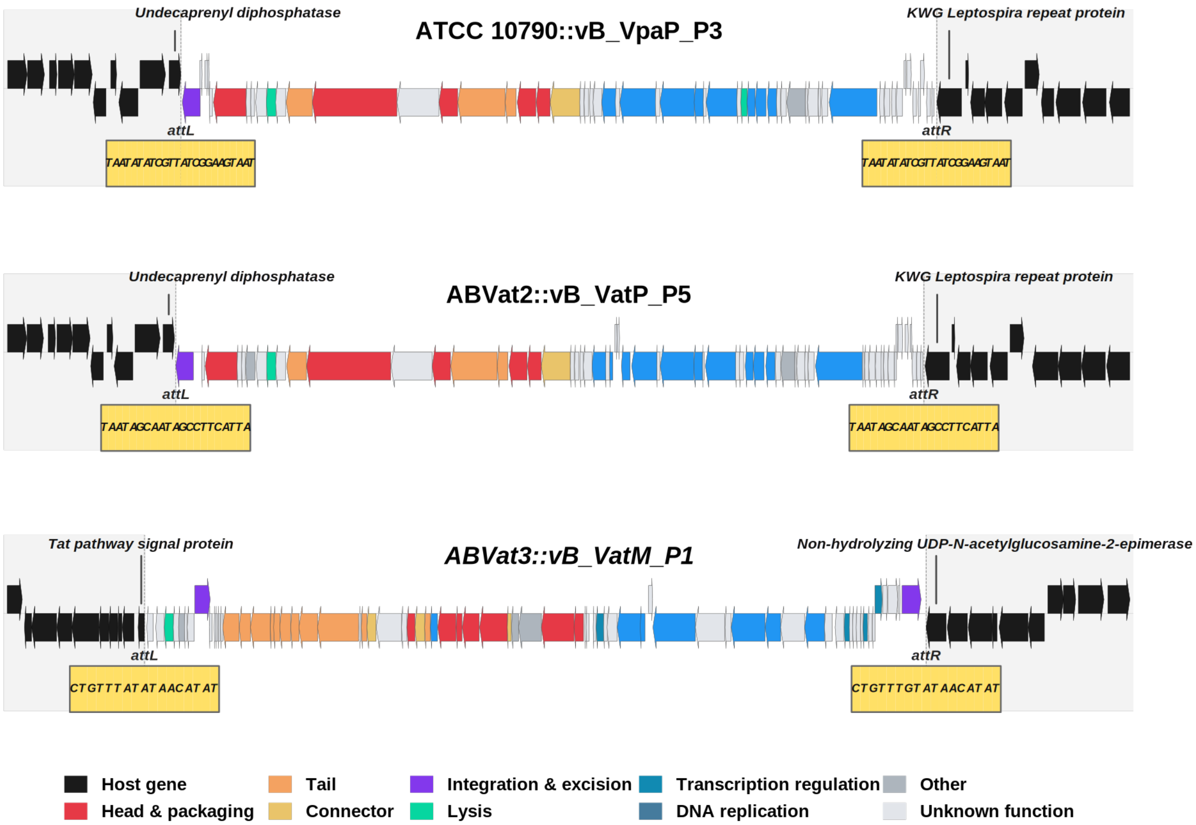


**Figure S4. Genomic integration sites of *Veillonella* phages.**
Schematic of the chromosomal integration locus for each of the three lysogens ATCC 10790::vB_VpaP_P3, ABVat2::vB_VatP_P5, and ABVat3::vB_VatM_P1. Genes are colored by functional category as in Fig. 2B. The attachment sequences attL/attR and the flanking host genes are labelled.

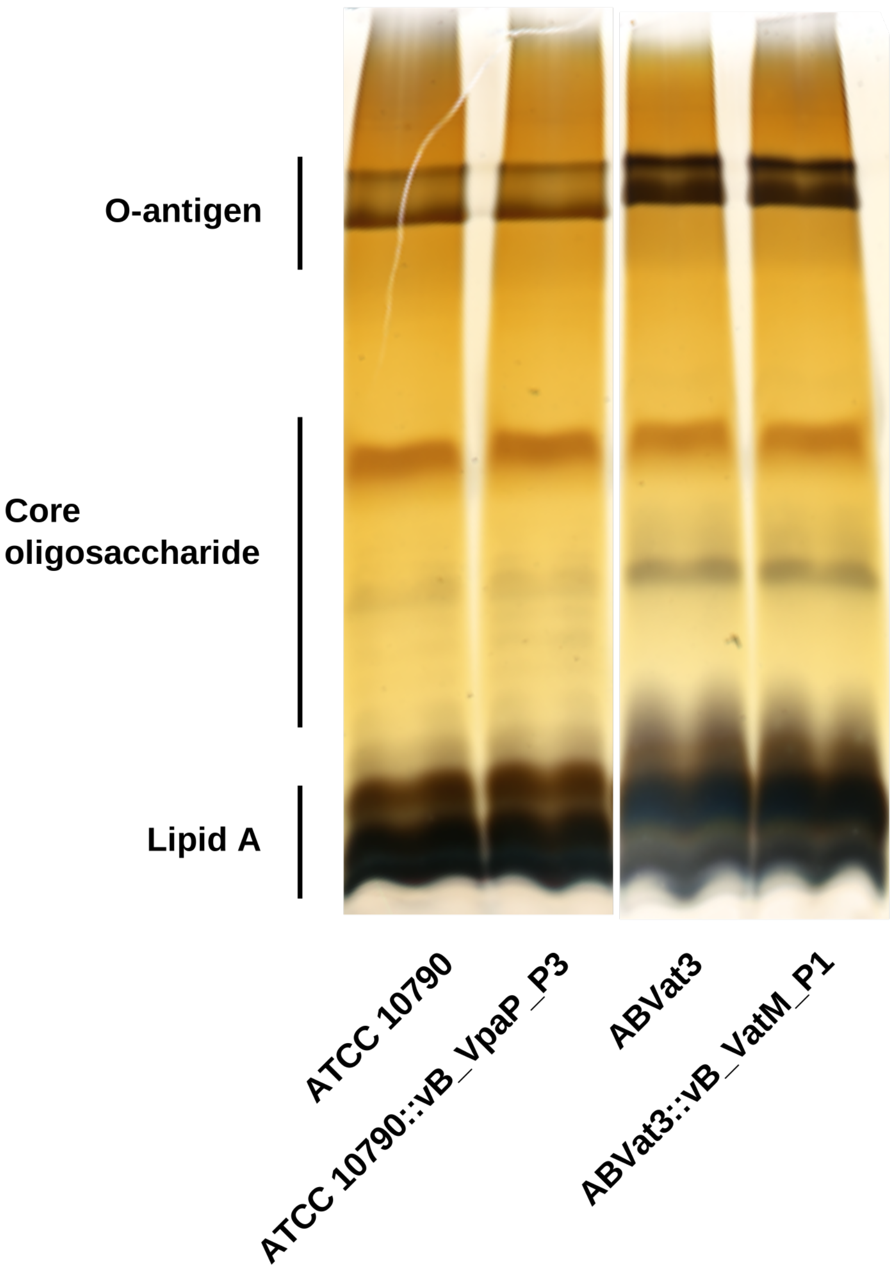


**Figure S5. Lipopolysaccharide (LPS) profiles of wild type and lysogenic *Veillonella* strains.**

Silver stained LPS profiles comparing ATCC 10790 and ABVat3 with their respective lysogens are shown. Locations of O-antigen, core oligosaccharide and lipid A are indicated, and they have no discernible differences in banding patterns.

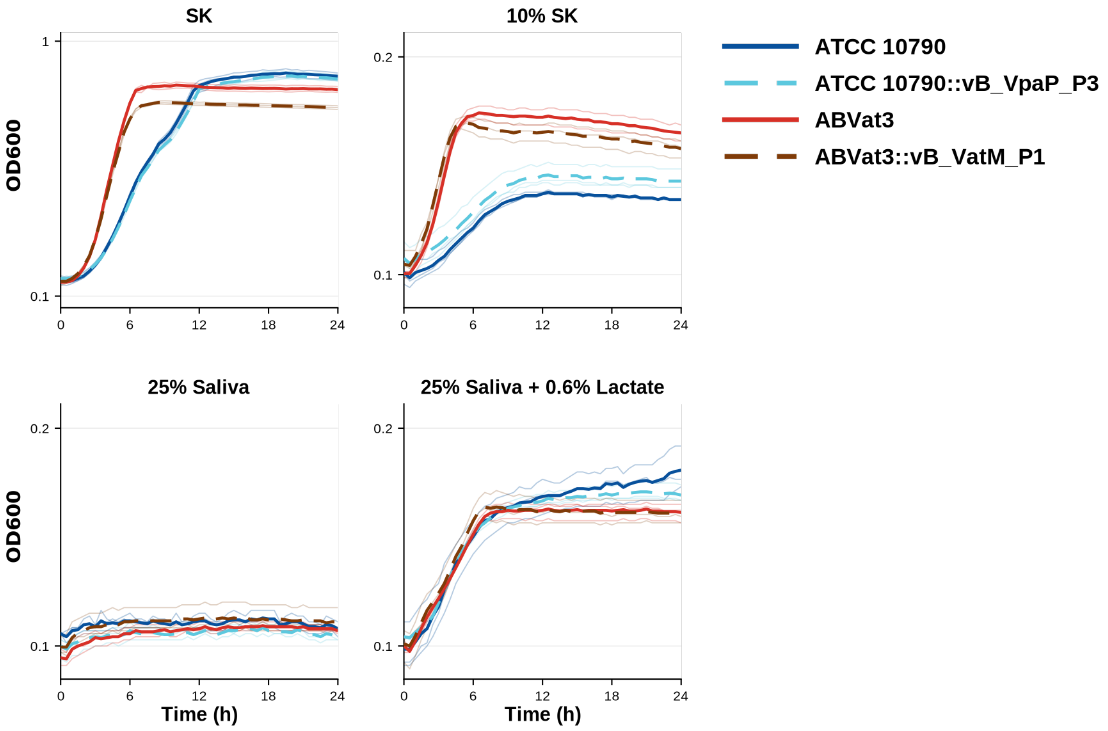


**Figure S6. Growth of wild-type and lysogenic strains across media conditions.**
Growth curves (OD600) of ATCC 10790, ABVat3 and their respective lysogens in SK, 10% SK, 25% saliva, and 25% saliva + 0.6% lactate over 24 hours. Lines represent means of n=3 biological replicates and shading indicates standard deviation. No consistent growth advantage was observed for the lysogen in any condition.

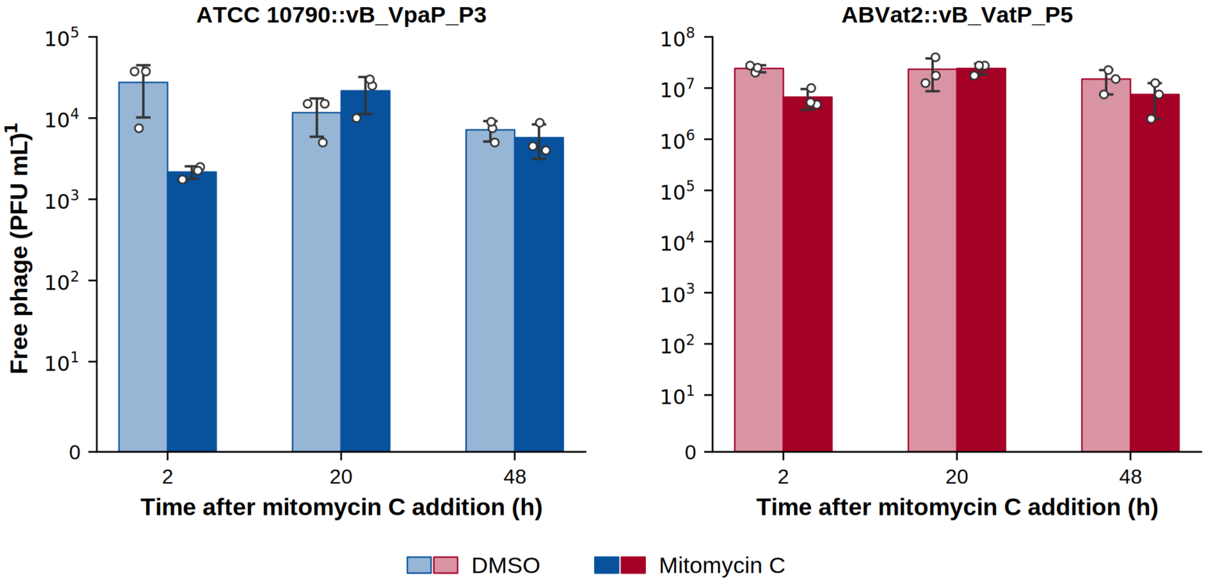


**Figure S7. Effect of mitomycin C on prophage induction.**

Bar plots showing free phage titers in the supernatants over time for lysogens ATCC 10790::vB_VpaP_P3 and ABVat2::vB_VatP_P5 following treatment with mitomycin C or a DMSO control. Points represent mean and SD of n=3 biological replicates.

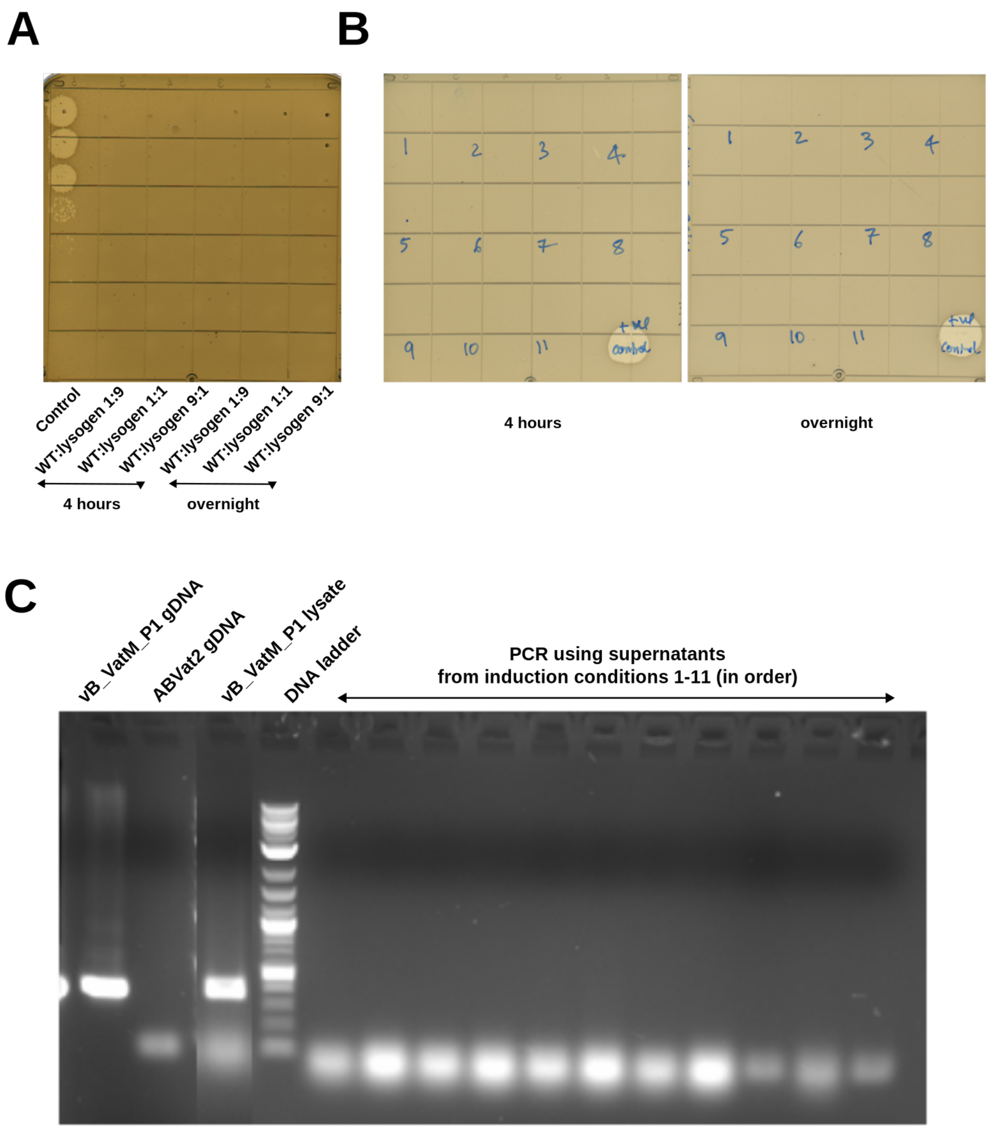


**Figure S8. Phage vB_VatM_P1 does not induce under coculture or diverse stress conditions.**

(A) Spot assay of filtered supernatants from ABVat3::vB_VatM_P1 cocultured with wild type ABVat3 at varying ratios for either 4 hours or overnight, spotted onto a *ABVat3* lawn to detect induced phage. A high titer lysate of vB_VatM_P1 served as a positive control. (B) Spot assay of filtered supernatants from ABVat3::vB_VatM_P1 following exposure to a panel of eleven stress conditions (**1**: 0.5 μg/mL mitomycin C, **2**: 2 μg/mL mitomycin C, **3**: 5 μg/mL mitomycin C, **4**: 1 mM $H_2O_2$, **5**: oxygen exposure, **6**: 42°C heat shock, **7**: 0°C cold shock, **8**: UV radiation, **9**: pH 4 acid, **10**: 10% SK starvation, **11**: SK + 10% saliva) and recovered for either 4 hours or overnight, spotted onto a wild-type ABVat3 lawn. A high titer lysate of vB_VatM_P1 served as a positive control. (C) PCR detection of vB_VatM_P1 in the same supernatants shown in (B), using primers targeting the phage integrase gene. Phage genomic DNA and heat treated lysate served as a positive control while host genomic DNA was a negative control.

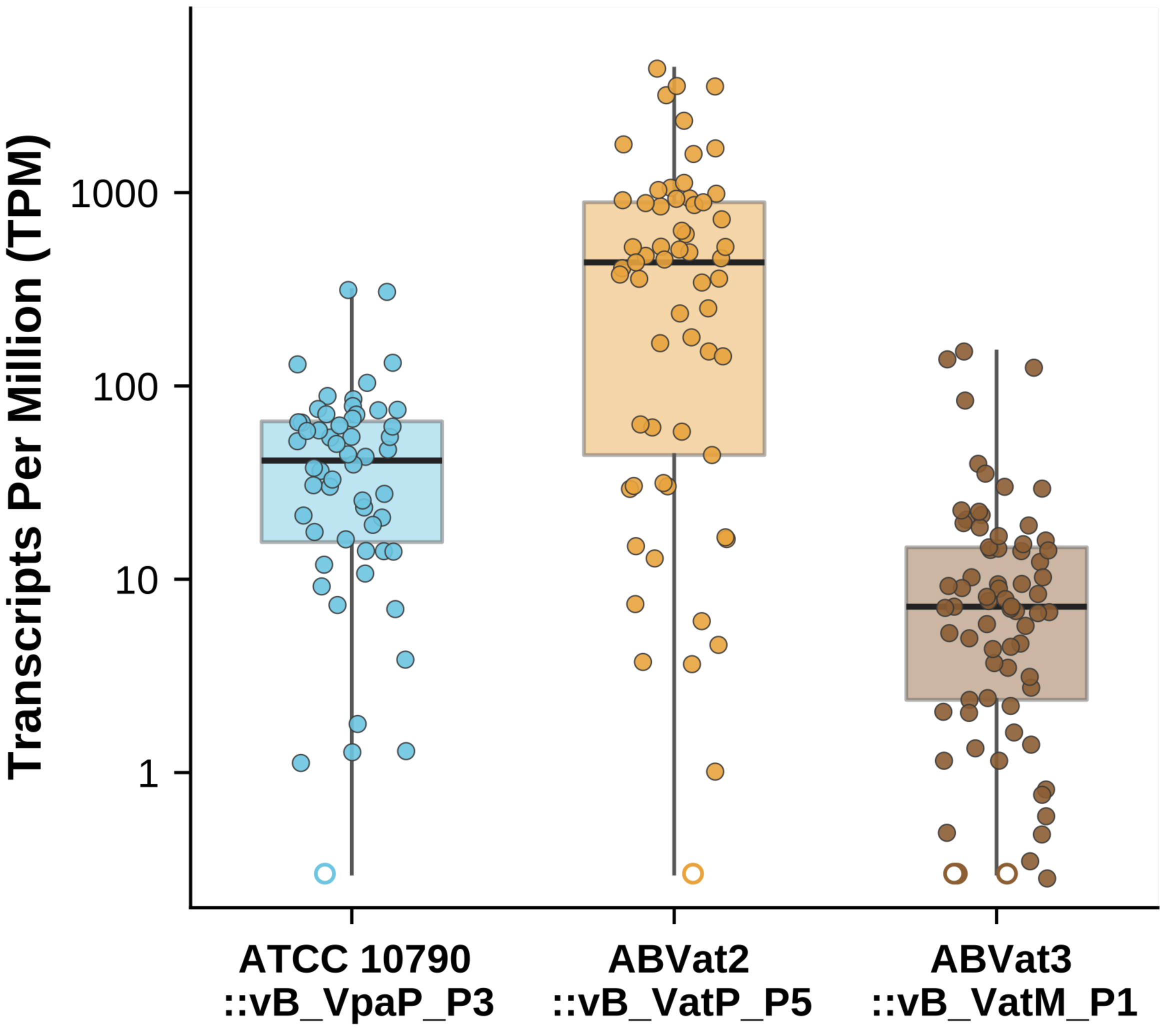


**Figure S9. Prophage gene expression differs by two orders of magnitude across lysogens.**

Transcript abundance of every prophage gene in three lysogens, from RNA-seq of exponentially growing cultures without induction. Each point is one gene and boxes give the median and interquartile range and whiskers span the full range. Open symbols at the axis floor are genes with no assigned fragments, which cannot be placed on a logarithmic axis.

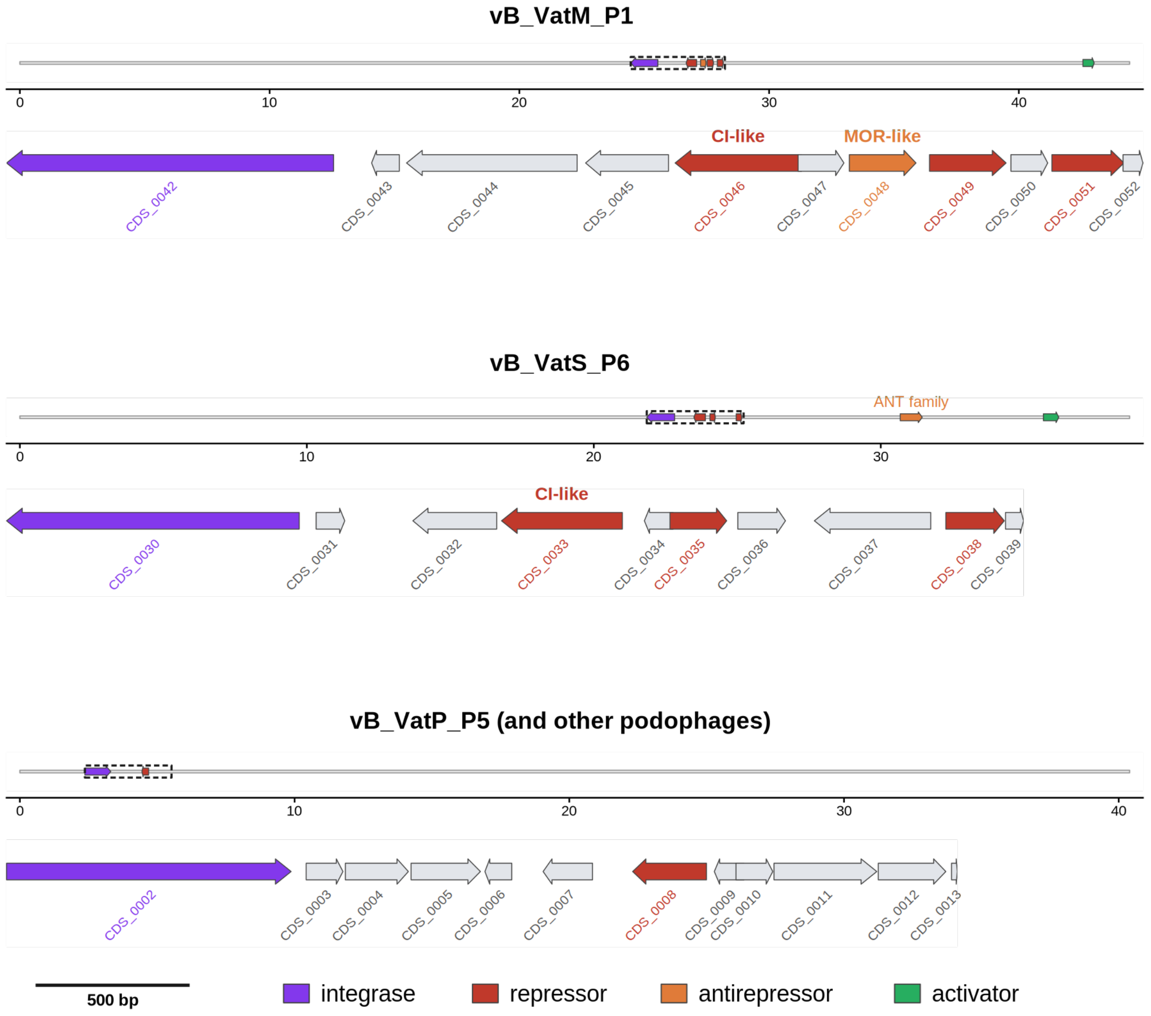


**Figure S10. Regulatory architecture of the lysis-lysogeny switch in novel *Veillonella* phages.**

Schematic showing lysis-lysogeny switches in the novel *Veillonella* phages. Upper track depicts whole genome with the integrase (purple), repressors (red), antirepressor (orange) and activator (green) in place and the switch region boxed and the lower track shows enlargement of that region drawn to a common scale (500 bp bar). Genes are labelled by locus tag and grey arrows have no predicted regulatory role. In vB_VatM_P1 the divergently oriented CDS_0046 and CDS_0048 carry HTH_3 (PF01381, E = $2.4 \times 10^{-20}$) and DUF739 (PF05339, E = $4.2 \times 10^{-8}$) and match the CI repressor and MOR antirepressor of Lactococcus phage TP901-1 (AAK38021.1, E = $4.1 \times 10^{-15}$; AAK38022.1, E = $2.5 \times 10^{-6}$). vB_VatS_P6 shares this arrangement, with CDS_0033 carrying HTH_3 and 41% identical over 129 aa to vB_VatM_P1_CDS_0046, and additionally encodes a distal ANT-family antirepressor (CDS_0047, PF03374, E = $3.5 \times 10^{-32}$). The podophages, represented by vB_VatP_P5, encode only a single small repressor adjacent to the integrase.
